# Plasmonic Enhancement of Protein Function

**DOI:** 10.1101/2024.02.07.579195

**Authors:** Marco Locarno, Qiangrui Dong, Xin Meng, Cristiano Glessi, Nynke Marije Hettema, Nidas Brandsma, Sebbe Blokhuizen, Alejandro Castañeda Garcia, Srividya Ganapathy, Marco Post, Thieme Schmidt, Lars van Roemburg, Bing Xu, Chun-Ting Cho, Liedewij Laan, Miao-Ping Chien, Daan Brinks

**Affiliations:** Department of Imaging Physics, Delft University of Technology, Delft, The Netherlands; Department of Bionanoscience, Delft University of Technology, Delft, The Netherlands; Department of Pediatrics and Cellular and Molecular Medicine, UCSD School of Medicine, San Diego, California, United States; Department of Radiation Science and Technology, Delft University of Technology, Delft, The Netherlands; Department of Molecular Genetics, Erasmus University Medical Center, Rotterdam, The Netherlands

## Abstract

Plasmonic nanoparticles are key components in nanophotonics^1–11^ and have applications in molecular detection and diagnostic platforms. ^12–14^ Coupling of dipoles to plasmonic antennas has allowed engineering of qualitatively altered behavior in isolated quantum systems, ^8,15^ but this promise has not been fulfilled in living systems, where the use of plasmonics is limited to tissue level applications of plasmonic particles as contrast agents^16^ or heat sources.^17^ Here we show that coupling to designed plasmonic nanoparticles can control the electrophysiological function of proteins in living cells. We designed nanostar geometries and achieved robust near-field coupling of these optimized nanoparticles to plasma membrane-localized Archaerhodopsin proteins. We enhanced the fluorescence of the coupled rhodopsins and increased their response speed to membrane voltage. We incorporated this plasmonic enhancement into a Markov chain photocycle model of the Archaerhodopsin mutant QuasAr6a, showing an increased fluorescence emission rate and manipulation of the protein dynamics through a combination of photocycle transition rate enhancements. These results represent the first scalable near-field coupling between plasmonic particles and fluorescent proteins in living cells. They show enhancement of protein function beyond what has been achievable through genetic engineering. This opens up a range of possibilities for engineering biological functionality through plasmonics.

---

The striking signal enhancements^7^ and altered quantum behavior^8^ that can be achieved by employing plasmonic nanoparticles in single-molecule fluorescence and SERS experiments^4^ depend on the exquisite near-field coupling between a plasmonic antenna and a molecular dipole of interest. These effects have been challenging to achieve in functional biological systems due to difficulties in the detection of small molecular signals over in vivo cellular backgrounds,^18,19^ accurate placement and orientation of nanoparticles for the required coupling,^20^ and robust creation of finely tuned nanoparticles.^21^ Nevertheless, if those issues can be overcome, plasmonics is a promising tool to achieve enhanced signals, increased resolution, and top-down control in the optical detection and manipulation of biological systems.^19^

Coupling to plasmonic particles can influence dipoles in two main ways, ^22^ both strongly distance-dependent.^20^ First, the plasmonic particle can localize an excitation field through resonance or lightning rod effect,^5^ leading locally to an enhanced electric field 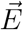 compared to a prior 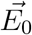, where the enhancement factor 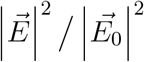 leads to a higher photon absorption per unit time. Second, the radiative decay rate of a dipole is given by Fermi’s golden rule ^2^ and can be manipulated by altering the Local Density of Optical States (LDOS) through careful positioning of dielectric or metallic elements in the presence of the dipole. ^22^ This effect can change the quantum yield of radiative transitions:

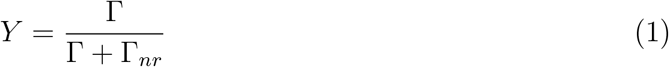

where *Y* is the radiative quantum yield of the transition, Γ is the radiative decay rate and Γ_*nr*_ is the non-radiative decay rate. **Eq. (1)** indicates that this can enhance low quantum yield processes (i.e. *Y* ≪ 1): in this case, Γ ≪ Γ_*nr*_. Plasmonic enhancement can increase Γ, leading theoretically to a *Y* approaching 1 if the effect is strong enough. ^22^

Our interest in this technology was fueled by two realizations regarding light-sensitive proteins: they often have a low fluorescence quantum yield,^23,24^ since they have typically evolved to use absorbed photons for performing a light-induced mechanical action;^25^ and this action typically incorporate a large number of transitions with excitable dipoles (the photocycle)^25,26^ which suggests not only fluorescence, but also protein dynamics, and with it, function, could be influenced through plasmonics.

We reasoned that judicious design and application of plasmonic particle geometry and spacing might allow us to achieve these effects in living biological systems. We found sparse reference to attempts at plasmonic enhancement of fluorescence of proteins, ^27,28^ one reference attempting fluorescence enhancement of fixed cells labeled with fluorescent dyes,^18^ and one in bacterial environments.^29^ However, we did not find attempts at this in electrophysiologically relevant conditions, let alone while incorporating influence on protein dynamics, and set out to test the feasibility of our idea.

## Results

### Particle design

For enhancement of transition rates, in a first-order approximation, the resonance spectrum of the plasmonic particles should have maximum overlap with the action spectrum (absorption/emission) of the dipole of interest.^22^ For biocompatibility, it is particularly interesting if this spectrum is red shifted, ensuring lower phototoxicity and easier light penetration, and tunable to the dipole of interest. ^18^ To achieve the best field enhancement, we first investigated the optimal particle geometry and material through Finite-Difference Time-Domain (FDTD) simulations of a series of nanospheres, nanorods and various types of spiked nanoparticles in biocompatible gold and silver. As expected from plasmonic theory^30^ gold spheres exhibit a suboptimal, broad peak in the range of interest (520 nm in water, **Figs. 1a**, **S1**). Gold nanorods are highly tunable, but the emergence of two resonant modes imposes the need for highly accurate orientation (**Fig. S2**), as it does for nanostars with a few sharp tips (**Figs. S3-S4**). However, highly spiked stars are tunable to the optimal wavelength in our range of interest (**Fig. 1a**) and their resonance is robust to rotation (**Fig. S5**). Furthermore, the localization of the field around nanospheres spans the entire surface of the sphere and reaches a squared field enhancement of the order of tens at best (**Fig. 1b**). In contrast, the local intensity of the electric fields around the sharp tips can be enhanced hundreds- or even thousands-fold (**Fig. 1c**).

**Figure 1:**
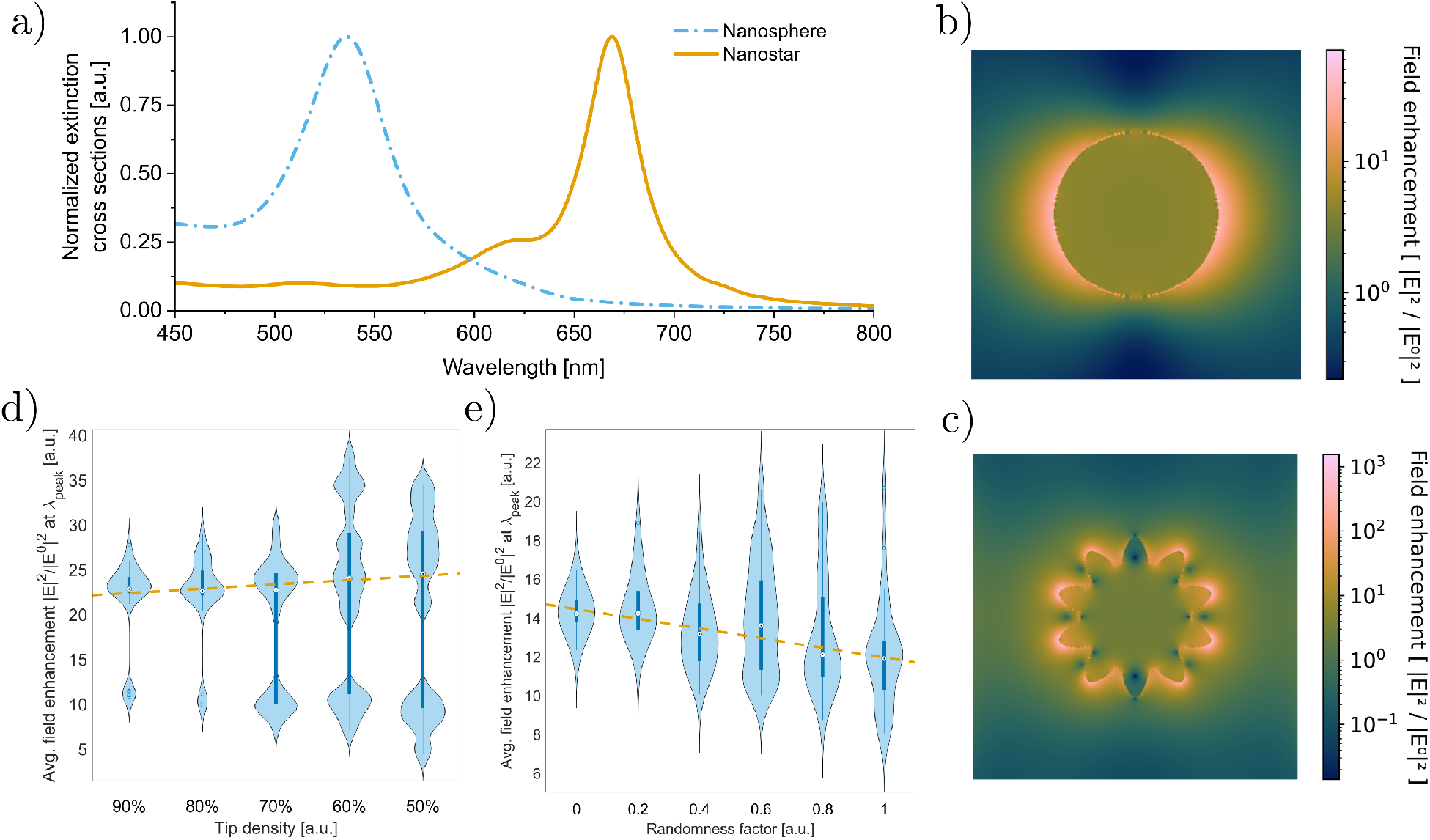
Gold nanostars are optimal candidates for in vivo plasmonic enhancement. **a**, FDTD simulated extinction cross sections of, respectively, a gold nanosphere and a gold nanostar in a watery environment. The simulated nanosphere has a radius of 25*nm*. The simulated nanostar has a core radius of 25*nm* and a tip length of 10*nm*. **b**, FDTD simulation of the field enhancement around a gold nanosphere in water at *λ* = 540*nm*. **c**, FDTD simulation of the field enhancement around a gold nanostar in water at *λ* = 650*nm*. The field is confined to a few nanometers around the nanostar spikes. **d**, FDTD simulations of gold nanostars with a core radius of 30*nm* and a tip length of 10*nm*, with different tip densities. For each simulation, the positions of the missing tips and the orientation of the nanostar were randomized. The individual results are shown in the Supplementary Information (**Fig. S6**). **e**, FDTD simulations of randomized gold nanostars from an initial ideal particle with a core radius of 30*nm* and a tip length of 10*nm*. The tip vertex position and nanostar orientation were randomized for each simulation. The individual results can be found in the Supplementary Information (**Fig. S7**).

The diffusive motion of nanoparticles in living biological samples might lead to changes in coupling distance and relative orientation to molecular dipoles, hampering plasmonic enhancement.^20^ Nanostars might also suffer from irregularities in their makeup, influencing their maximal enhancement.^31^ We quantified both effects and found that the enhancement by nanostars was robust against these sources of noise (**Figs. 1d,e**, individual simulations in **Figs. S6-S7**).

### Colloidal growth of nanostars

Gold is a common metal for plasmonic applications, characterized by superior biocompatibility and low cytotoxicity,^32^ and was, therefore, the material of choice used throughout this study. To synthesize the gold nanostars, we adapted a colloidal, seed-mediated growth protocol^31^ that allowed high control and tunability of the geometry. We synthesized a series of nanostars under varying seed and growth conditions (**Methods section**). We characterized the synthesized colloids through Energy Dispersive X-ray Spectroscopy (EDX, **Fig. S8**) and Ultraviolet-Visible light (UV-Vis) spectroscopy. The optimal sample (molar ratio *R* = 30) was selected based on the alignment of the absorbance peak to the emission peak of Archon1^33^ (**Fig. S9a**). We attribute a minor mismatch to a red shift induced by polyvinylpyrrolidone (PVP) as a capping agent (**Eq. (S4)**).

We performed quality control by immobilizing the gold nanostars on a silicon substrate with 3-mercaptopropyl trimethoxysilane (MPTMS) (**Fig. S9b**), following a custom protocol (**Methods section**), to examine them under a scanning electron microscope (SEM). The monodisperse nanostars had an average core radius of *r* = 23 ± 5 nm (mean ± SD, **Fig. S9c**) and an average tip length *l* = 12 ± 3 nm, (mean ± SD, **Fig. S9c**) in agreement with the simulations. They were generally well separated (**Fig. S9d**), though occasional clustering in micrometer-sized aggregates was visible (**Fig. S9e**), possibly affecting the measured plasmonic resonance.^12^

### Calibration using Cyanine-5

Next, we optimized a protocol to couple plasmonic particles to fluorescent dipoles in wet lab conditions. We chose Cyanine-5 (Cy5) as a model molecule to enhance the fluorescence because of its emission peak position and moderate quantum yield (∼30%).^34,35^

We hypothesized that Biotin-Streptavidin binding would provide a suitable way to link nanostars to fluorescent emitters under physiological conditions, and compared the fluorescence readout of Streptavidin-Cy5 (Strep-Cy5), a mixture of Strep-Cy5 and citrate-capped nanostars, and Strep-Cy5 bound to biotin-coated nanostars (**Fig. 2a**) However, we found that the presence of nanostars reduced the fluorescence signal by up to -68 ± 16% (mean ± error, propagated from SD, **Fig. 2b**). We attribute this to a combination of the increased optical density in nanostar-containing samples,^36^ and fluorescence quenching caused by non-radiative energy transfer from the fluorophore to the nanostar.^1^ Streptavidin, the largest molecule in the binding complex, is around 5 nm,^37^ keeping the fluorophore within the quenching range of the nanostar. ^1^

**Figure 2:**
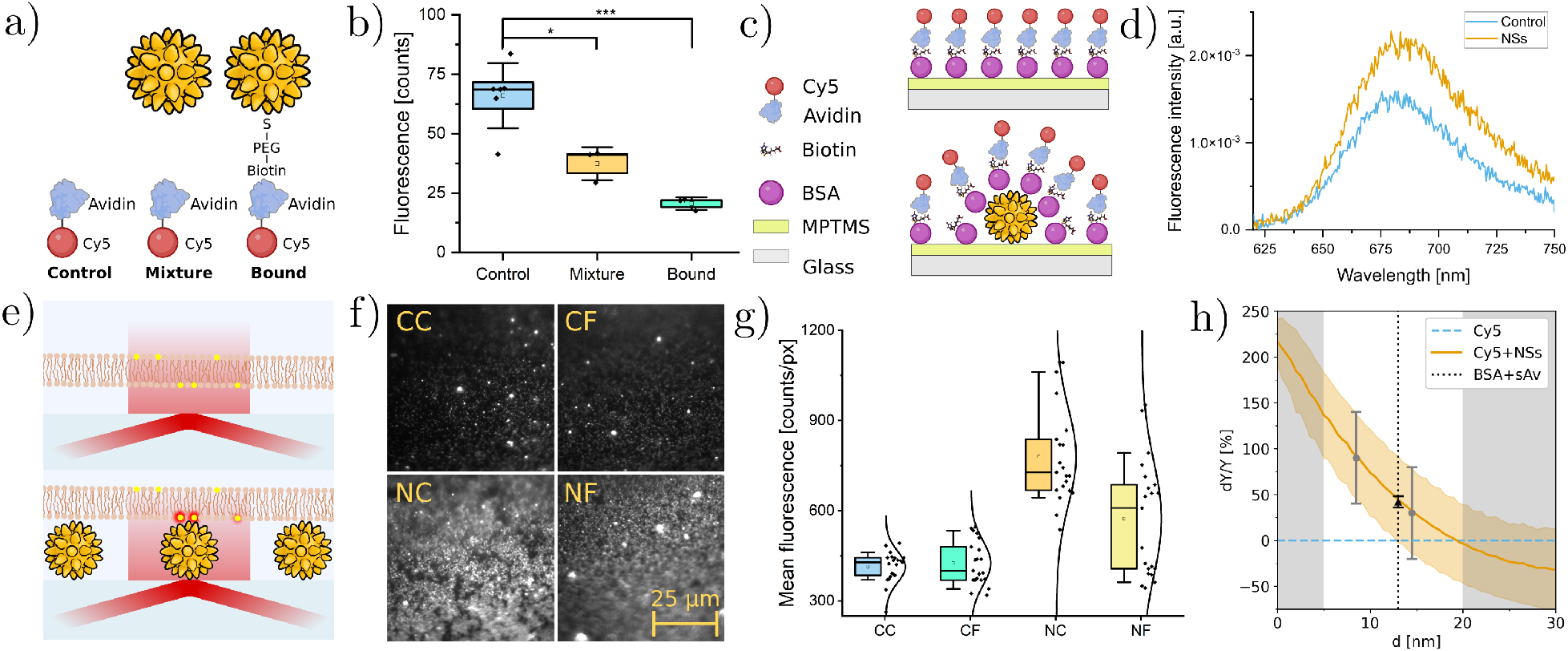
Control over nanostar-fluorophore coupling allows tunable plasmonic enhancement. **a**, Illustration of fluorescence assays in solution. control: Cy5 labelled with streptavidin; mixture: control sample mixed with citrate-capped commercial nanostars; bound: Strep-Cy5 bound to biotin PEGylated gold nanostars. Biotin and streptavidin bind, ensuring a fixed distance between the nanostar and the fluorophore. Not to scale. **b**, Results from fluorescence spectroscopy at peak wavelength (666 nm). Boxes represent the standard errors, whiskers the SD, squares the means and mid-lines the medians. **c**, Illustration of fluorescence assay on glass substrate. **d**, Results from fluorescence spectroscopy in glass dishes. **e**, Illustration of TIRF microscopy of a fluorescently-labelled artificial membrane without (top) and with nanostars (bottom). Not to scale. **f**, Median TIRF images of the artificial membranes. Represented with the same brightness and contrast. **g**, Cross comparison of the fluorescence in DOPC lipid bilayers labelled with Cy5 dye, without Fibronecton or Nanostars (CC, 410 ± 50 counts/px), with Fibronectin (CF, 430 ± 70 counts/px), with Nanostars (NC, 780 ± 160 counts/px) or with Nanostars and Fibronectin (NF, 570 ± 190 counts/px). Boxes represent the 25-75% range, whiskers the 10-90% range, squares the means and mid-lines the medians. **h**, Predicted mean enhancement of Cy5 (orange, solid line) within the range of uncertainty (orange band, propagated from SD), based on simulation, compared to the natural *Y* of Cy5 (light blue, dashed line). Data points represent the measured enhancement in panel d (mean as the black triangle, error bar propagated from noise SD) and in panel g (means as the gray dots, error bars propagated from SD). Distances for measurements from g are determined by using the curve as a plasmonic ruler.

This experiment provided a baseline distance measurement, and we set out to increase the distance between the nanostars and the emitters. Using MPTMS, we immobilized gold nanostars functionalized with a complex of Bovine Serum Albumin (BSA) and biotin on glass, then added Strep-Cy5 (**Fig. 2c**). Fluorescence spectroscopy showed a net 42 ± 6% increase in the fluorescence intensity of Cy5 (peak value ± error, propagated from noise SD, **Fig. 2d**), in agreement with the theoretical value (**Eq. (S7)**) considering the combined lengths of streptavidin (5 nm) ^37^ and BSA (8 nm)^38^ as a spacer larger than the quenching range,^1^ the already significant quantum yield of Cy5,^34,35^ and the estimated average field enhancement of a factor 1.78 from simulations (**Fig. S10**).

To apply plasmonic enhancement in living cells, this effect needs to be compatible with cell culturing conditions.^18^ For ease of coupling nanoparticles, we focus for now on proteins that are embedded in the lipid bilayer of the cell membrane.^25^ Cells are typically attached to glass substrates using an extracellular matrix protein, like fibronectin.^39^ Given the sensitivity of the enhancement to the dielectric environment, we wondered whether the lipid bilayer or fibronectin would influence the enhancement. We immobilized nanostars on glass and used Total Internal Reflection Fluorescence (TIRF) microscopy to investigate the effect of an artificial lipid bilayer and a standard fibronectin layer (**Methods section, Fig. 2e**). In this in vitro experiment, we positioned lipid bilayers labelled with Cy5 on the sample and imaged the fluorescence with and without nanostars, and with and without fibronectin (**Fig. 2f**). We measured a fluorescence enhancement of 90 ± 50% (mean ± error, propagated from SD) associated with the direct coupling between Cy5 and nanostars, while the presence of the fibronectin layer reduced the effect to 30 ± 50% enhancement (mean ± error, propagated from SD), probably by acting as a spacer (**Fig. 2g**).^1^ In combination with the Strep-Cy5 results (**Fig. 2d**), we estimate the distance between the Cy5 molecules and the nanostars to be 14.5 nm, respectively 8.5 nm with and without fibronectin (**Fig. 2h**), for an effective 6 nm thickness of the fibronectin layer.

### Plasmonic enhancement in living cells

To test our system in living cells, we chose Archaerhodopsin as a model light-sensitive protein. Several mutations of this protein exist (Quasar1-6a,b;^24,40,41^ Archon1,2;^42^ NovArch^43^), which are all known for low quantum yield of fluorescence. Their emission spectrum, which was measured to be between 640 and 800 nm,^23,33^ is in an interesting range for bioimaging applications. Moreover, Archaerhodopsin3 is a voltage-sensitive proton pump,^23,24,41,44^ and all of its mutants have a photocycle and fluorescence that depend on the voltage across the cell membrane in which the protein is embedded, making it an interesting proof of principle model for manipulation of protein function using plasmonics.

Having shown that cell membrane embedding and fibronectin application do not hinder plasmonic enhancement of fluorophores, we performed a simple addition of colloidal nanostars to HEK293T cells expressing either QuasAr1 (**Fig. 3a**) or QuasAr6a (**Fig. 3b**).

**Figure 3:**
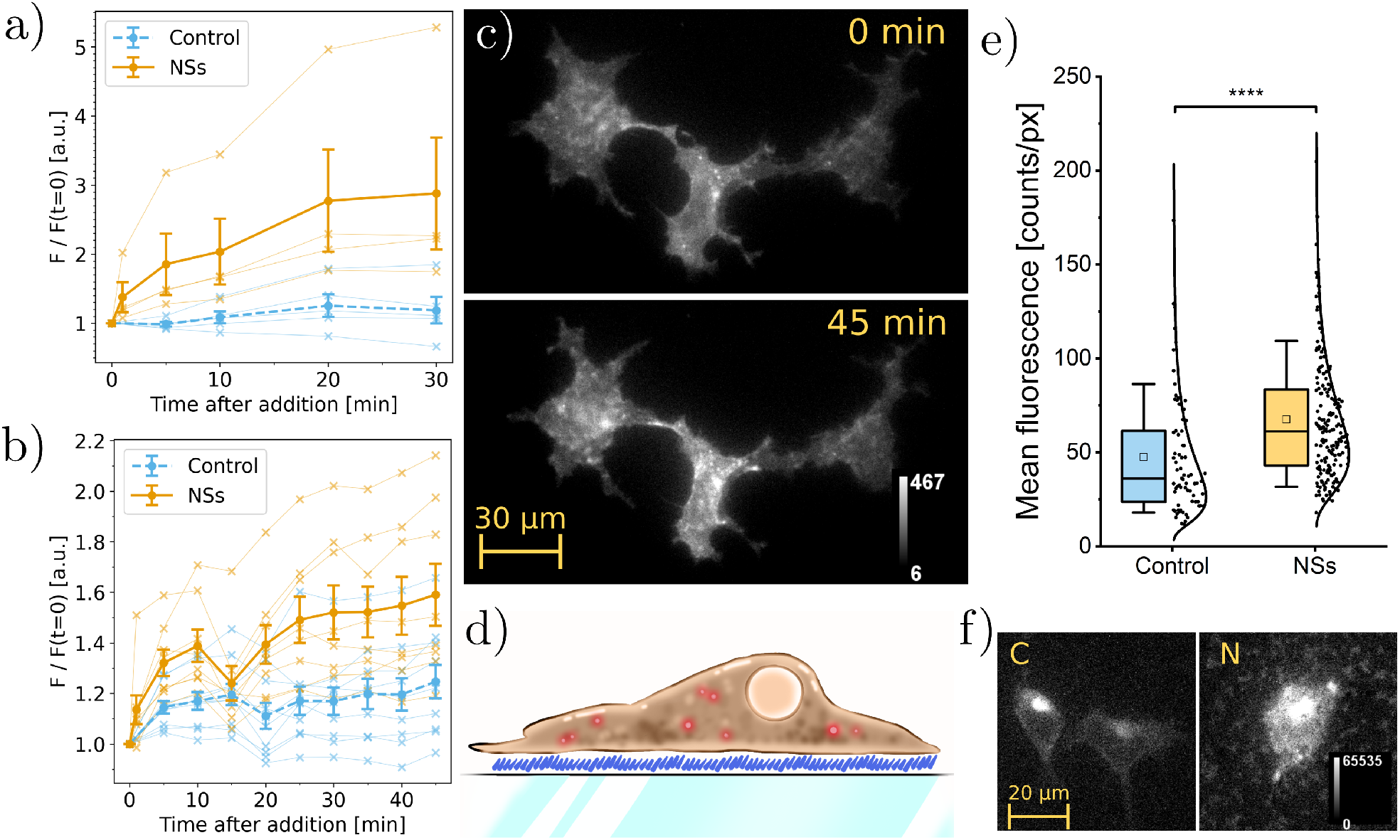
Gold nanostars enable plasmonic enhancement of QuasAr fluorescence in living cells. **a**, Mean fluorescence of cells expressing QuasAr1, normalized to the first frame, after the addition 400*µl* of nanostars solution (NSs), compared to no addition (control). Error bars correspond to the standard error. Over time, the nanostars gradually diffuse and sediment on the cells, enhancing the fluorescence. Crosses indicate individual measurements, filled circles averages across all trials. **b**, Mean fluorescence of cells expressing QuasAr6a, normalized to the first frame, after the addition 400*µl* of nanostars solution (NSs), compared to no addition (control). Error bars correspond to the standard error. Crosses indicate individual measurements, and filled circles averages across all trials. **c**, Example of fluorescent cells in the QuasAr6a test sample before and after the addition of nanostars. **d**, Illustration of a fluorescent HEK293T cell plated on fibronectin, layered above a glass dish covered in nanostars (in blue). **e**, Screening results show a statistical difference between a sample without (control) and with nanostars (NSs). Boxes represent the 25-75% range, whiskers the 10-90% range, squares the means and mid-lines the medians. Uncertainty propagated from the standard error. **f**, Comparison of the median cell in the control screening (left) and in the nanostars sample (right). Individual nanostars responsible for enhancement are too small to be distinguished but nanostar aggregates are visible either as dim autofluorescence outside the cell, or spot-like shadows under the cell.

We noticed an enhancement in the fluorescence of individual cells over time (**Fig. 3c**). We quantified the change in fluorescence of the cell population over time after the addition of the nanostars and noticed an increase of fluorescence to a factor 140 ± 80% (QuasAr1, mean ± error, propagated from SE, **Fig. 3a**) and 28 ± 12% (QuasAr6a, mean ± error, propagated from the SE, **Fig. 3b**) respectively.

We tested the difference between this simple application of nanostars and embedding them in the fibronectin layer. We immobilized gold nanostars on glass, coated the surface with fibronectin, and plated HEK293T cells expressing QuasAr6a (**Fig. 3d**). We screened the fluorescence at the bottom surface of the cell population (**Fig. 3e**) and achieved a fluorescence enhancement of 69 ± 19% (median ± error, propagated from the SE), confirmed by the fluorescence brightness of the median cell in a control, respectively enhanced sample (**Fig. 3f**). The difference in enhancement of fluorescence between the nanostars applied to the top surface (28 ± 12%, **Fig. 3b**) to those embedded in the fibronectin adhesion layer (69 ± 19%, **Fig. 3e**) indicates that the glycocalyx^45,46^ surrounding the cells acts effectively as a spacer between the fluorophores and the nanoparticles applied to the top surface of the cells. The glycocalyx is thinner at the bottom of the cells because the formation of integrin-fibronectin bonds induces mechanical strain. ^47^ We estimated an effective distance from QuasAr6a of 13.5 nm for the nanostars embedded in the fibronectin layer and 16 nm for the nanostars deposited above the cells; we estimate the glycocalyx to cause an average of 8.5 nm spacing difference (**Eq. (S15)**).

### Plasmonic enhancement of protein function

Fluorescence enhancement of dim genetically encoded voltage indicators like QuasArs is one possible application of coupling plasmonic antennas to proteins in living cell environments. Given that the kinetics of GEVIs are determined by a photocycle with potential absorption and emission events at multiple stages,^25,44,48^ we were interested in whether coupling to plasmonic antennas might also be used to enhance the protein function. We performed a voltage clamp experiment on HEK293T cells expressing QuasAr6a plated on fibronectin over immobilized gold nanostars to test this. We varied the voltage across the cell membrane in steps from -70 to +30 mV and recorded the fluorescence response without (**Fig. 4a**) and with nanostars in the fibronectin layer (**Fig. 4b**). We noticed a decrease in voltage sensitivity from a ΔF/F of 0.37 ± 0.04 to 0.11 ± 0.04 (mean ± SEM, **Fig. 4c**), mostly caused by the increase in baseline fluorescence, rather than a change in the absolute response to voltage.

**Figure 4:**
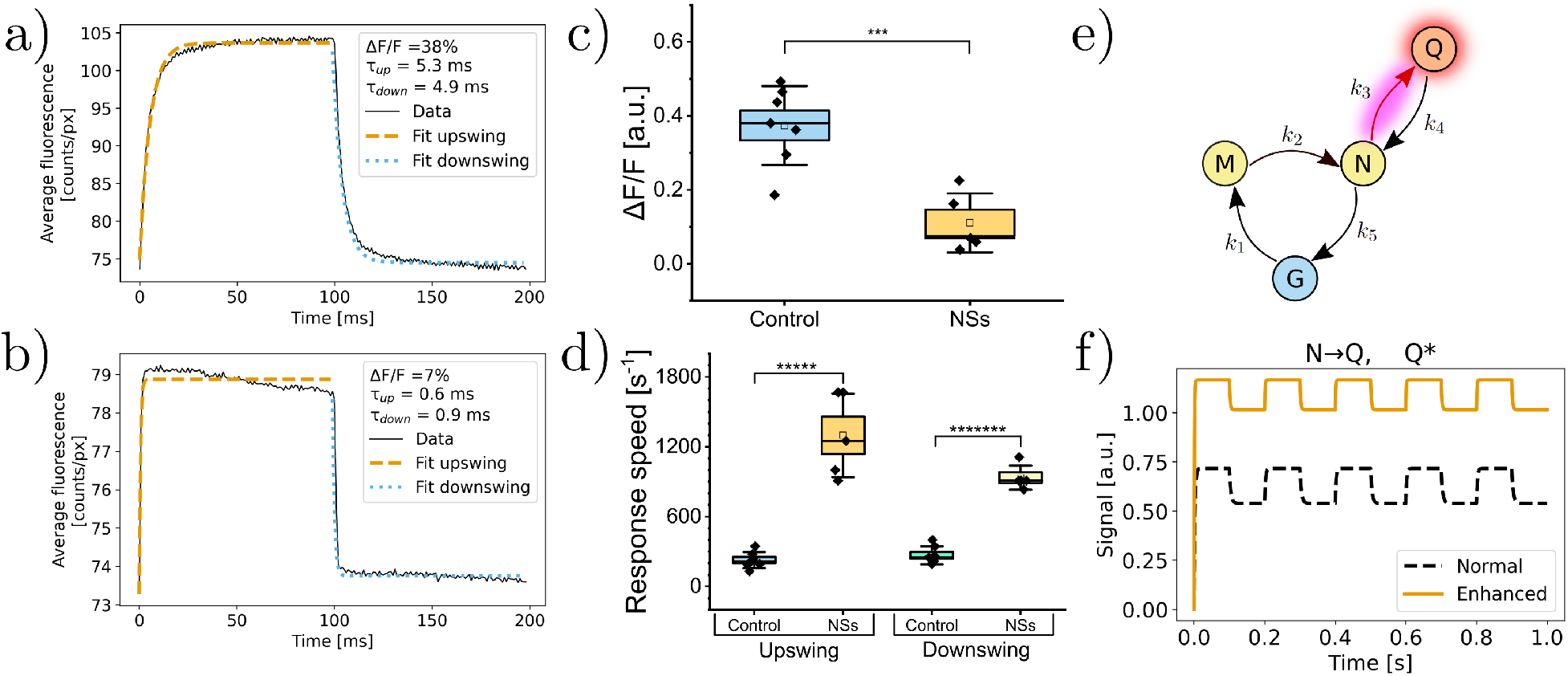
Plasmonic enhancement by gold nanostars manipulates the photocycle of QuasAr6a and causes enhanced brightness, diminished voltage sensitivity, and faster kinetics. **a**, Average trace of the median control cell under the influence of a square wave in membrane potential (−70 mV to +30 mV). **b**, Like a, but in the presence of nanostars. **c**, Voltage sensitivity of QuasAr6a without (control) and with nanostars (NSs). Boxes represent the standard errors, whiskers the standard deviation, squares the means and mid-lines the medians. The mean sensitivity of the control is compatible with the value reported in the literature 0.43 ± 0.04. ^41^ The mean sensitivity decreases by 71% compared to the control. **d**, Response speed of QuasAr6a without (control) and with nanostars (NSs). Boxes represent the standard errors, whiskers the standard deviation, squares the means and mid-lines the medians. Both the upswing and the downswing transition become faster when the fluorophores are coupled with the nanostars. **e**, Proposed photocycle for QuasAr6a; fluorescence is thought to come from the Q-state, *k*_2_ is voltage-dependent and *k*_3_ is photoactivated. **f**, Numerically simulated traces for the apparent occupancy of the Q-state of QuasAr6a under the influence of a square wave in membrane potential (−70 mV to +30 mV). Plasmonic enhancement is assumed to enhance *k*_3_ and the quantum yield of the Q-state.

However, we noticed a strong increase in response speed from *k*_*up*_ = 230 ± 70 s^−1^ to 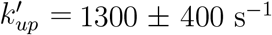 and *k*_*down*_ = 270 ± 80 s^−1^ to 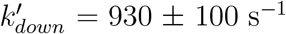 (mean ± SD, **Fig. 4d**)

Following the template photocycle for Arch, ^44^ we modeled a phenomenological photocycle for QuasAr6a as a continuous-time finite-state Markov chain (**Fig. 4e**).^49^ Here, the M→N transition is voltage sensitive, the N→Q transition is photosensitive, and Q is the excitable fluorescent state. We modeled fluorescence as proportional to the Q-state population multiplied by field enhancement and fitted our data to models with varying transition rates and field enhancement. We found the best fit to the data, including enhanced brightness, decreased voltage sensitivity, and faster kinetics, for a model incorporating *k*_3_ = 443 s^−1^ without plasmonic enhancement and *k* = 1184 s^−1^ and 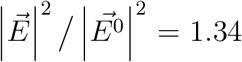 with plasmonic coupling (**Fig. 4f**, **Fig. S11**, complete parameters and results available in **Tables S1-S2**). Enhancement of only the quantum yield evidently does not explain the change in kinetics (**Fig. S11a**), and enhancement of only *k*_3_ leads to a worse fit of the data (**Fig. S11b**). Based on this model, we interpret the increased fluorescence and response speed to voltage as caused by differential enhancement of absorption of the photon promoting the N→Q transition and quantum yield of the Q-state fluorescence, tunable by the resonance spectrum of the plasmonic antenna.

## Discussion

This study demonstrates the use of plasmonic enhancement to engineer the function of proteins in living cells, which has implications for biosensing and functional bioimaging applications. We enhanced the fluorescence and voltage response speed of membrane-localized Archaerhodopsin proteins. The emergence of these noticeable effects, even without precise control over the coupling distance, suggests a robust effect that could potentially be applied in animal models. Moreover, because plasmonic enhancement is a fast physical phenomenon lasting tens of femtoseconds,^9^ it would be compatible with multiphoton excitation or other nonlinear imaging methods.

Archaerhodopsins are natural proton pumps; ^50^ the QuasAr and Archon mutants have been created to break the proton pumping function so that these proteins can be used as GEVIs.^23,24^ These proteins have gone through years of protein engineering efforts to increase their functionality.^23,24,33,41,43,44^ Despite this, none of the evolved versions have succeeded in creating a sensor that responds to membrane voltage faster than the original Archaerhodopsin3. The coupling to a plasmonic nanoantenna created a hybrid construct that enhanced the core function of the protein, its response to membrane voltage, to levels never achieved before by genetic manipulation, showing the promise of top-down control of protein function using plasmonics. Protein evolution of GEVIs often leads to increases in brightness at the expense of decreases in the speed of response to voltage changes. ^26^ Plasmonic enhancement shows a way to achieve both effects simultaneously. We hypothesize that a controlled trade-off between the enhancement of absorption (*k*_3_) and emission (*Y*) can be used to fine-tune voltage sensitivity, speed, and brightness of GEVIs.

The applicability of this concept is enhanced by well-established methods to link (gold) nanoparticles to biomolecules, ^17^ which would allow for even more precise control of distance and orientation of particles. The dense-star geometry shown in this work is also, in this combination, the critical shape for a plasmonic antenna, due to the robustness of its coupling against positional and orientational noise.

In conclusion, we showed the design and creation of gold nanostars for use as plasmonic antennas in live cell environments and achieved enhancement of protein function of mutated Archaerhodopsin proteins. We enhanced fluorescence and response speed to voltage of QuasAr6a by tunable coupling to plasmonic nanostars in live cells. This work creates possibilities for application of nanoantennas in bioengineering, particularly in engineering protein functionality, from light harvesting to sensing and top-down influencing of signaling cascades.

## Methods

### FDTD simulations

Finite-difference time-domain (FDTD) numerical simulations were conducted using Lumerical’s FDTD Solutions, with scripts available on GitHub.^51^ Simulations of gold nanospheres involved varying the radius *R* of a gold sphere submerged in water, with a focus on assessing the maximum field enhancement |*E*|^2^*/*|*E*^0^|^2^ within a small cubic volume (Δ*x* = Δ*y* = Δ*z* = *R/*2) located 4 nm from the surface. Simulations covered a range of radii (*R*_min_ = 6 nm to *R*_max_ = 76 nm) with 2 nm steps.

Gold nanostar models were crafted based on SEM micrographs of synthesized nanoparticles. To generate these models, Blender software was employed to create complex meshes. The starting point for this modelling was an icosahedron shape, with its vertices extended to mimic the tip arrangement found on real nanostars. To ensure smoother surfaces, a Subdivision Surface Modifier was applied. During these simulations, a mesh size of 35 Meshes Per Radius (MPR) was consistently used. For benchmarking, a model nanostar with a core radius *R* of 25 nm and a tip length *L*_tip_ of 10 nm was adopted. For the simulations shown in **Figs. 1d** and **1e**, a core radius of 30 nm was used. Simulations explored variations in tip density and randomness to understand their implications for field enhancement. These variations included the random omission of tips and the random displacement of tip vertices, quantified using a ”randomness factor” in Blender ranging from 0 to 1. Each group consisted of 25 independent simulations. Since the sharp tips simulated in **Figs. 1a, 1c**, and **1d** generated unphysical fields when randomized in Blender, we introduced an adaptation to their sharpness to prevent unrealistic stretching and overly sharp edges. All scripts for Lumerical and Blender can be found on GitHub.^51^

### Colloidal synthesis

In the presence of a high concentration of polyvinylpyrrolidone (PVP) dissolved in dimethylformamide (DMF), highly branched nanostars can be obtained efficiently at room temperature by optimizing the molar ratio *R*=[HAuCl_4_]/[seeds].^21^ Gold tips can grow on pre-formed gold seeds thanks to the reducing power of DMF and the excess of PVP. The protocol for making PVP-coated gold nanostars was adapted from the seed-mediated synthesis proposed by Barbosa et al.^31^ For each trial, 15 mL of 10 mM PVP (Sigma-Aldrich, PVP10) solubilized in DMF (Sigma-Aldrich, D4551) were transferred into a 25 mL conical flask under continuous magnetic stirring. Then, 82 *µ*L of 50 mM HAuCL_4_ aqueous solution were added. Within 30 s, a predetermined volume of gold spherical seeds (in-house synthesis based on Barbosa et al.,^31^ estimated concentration 0.5 mM, 15 nm in diameter) was added. The volume of pre-formed seeds was determined based on the formula 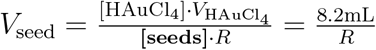 where *R* is the desired molar ratio. After 15 minutes under continuous stirring, the samples were spun at 4000 rpm for 90 minutes and the supernatant was removed to prevent the undesired formation of suboptimal nanoparticles. The nanostars were re-dispersed in MilliQ water and vortexed for a couple of seconds. This last purification step was carried out twice to minimize the cytotoxic effect of DMF and to reduce the amount of PVP in the solution. The optimal sample used throughout the manuscript corresponds to a molar ratio of 30 (273 *µ*L of seeds).

### UV/Visible spectroscopy of gold nanostars colloid

The absorbance spectra were measured using a compact Visible-NIR light source (ecoVIS by Ocean Optics) together with a compact STS spectrometer (STS-VIS Microspectrometer, 350 – 800 nm, by Ocean Insight). Absorbance was measured using an integration time of 5.1 ms after background subtraction and intensity calibration, and averaged 1000 times.

### MPTMS surface coating and nanoparticle immobilization

For the preparation of the MPTMS-coated dishes, a 10% MPTMS solution was prepared as follows: MPTMS (2.50 mL, Sigma-Aldrich, 175617) was combined with MilliQ water (2.50 mL), acetic acid (1.67 mL, Sigma-Aldrich, ARK2183), and pure ethanol (18.3 mL, Sigma-Aldrich, EMSURE 1.00983.1000). Prior to use, glass bottom dishes (Cellvis, D35-141.5-N) and fragments of silicon wafers were cleaned using oxygen plasma for 1 to 2 minutes, and all subsequent reactions were completed within a 30-minute timeframe. To prepare the substrates for cellular imaging, approximately 150 *µ*L of the MPTMS solution was gently dispensed into the recessed wells of the imaging dishes. These dishes were then carefully transferred to a large glass beaker and purged with nitrogen for 30 minutes (or until dry). Any residual MPTMS was aspirated. The dishes were subsequently subjected to a series of rinsing steps, including one wash with 150 *µ*L of pure ethanol in the recessed well, one wash with 2 mL of pure ethanol across the entire dish, and a final rinse with 2 mL of MilliQ water across the entire dish. To ensure dryness, the dishes were placed in a vacuum oven at 65°C for 2 hours. 200 *µ*L of gold nanostars were uniformly added to cover each recessed well. The dishes were then incubated overnight in a 37°C incubator, ensuring their cleanliness and contamination-free environment. Subsequently, the nanostar-coated substrates were aspirated, subjected to two 2 mL MilliQ water washes, and dried to near dryness. They were stored in the incubator until their intended use. For silicon substrates, the protocol remained unchanged, with volumes adjusted proportionally to the chip’s surface area.

### SEM/EDX characterization

SEM imaging was performed in a Thermo Fisher Scientific (TFS) Nova 600i Nano Lab dualbeam system with a primary beam energy of 5 kV and nominal current of 0.27 nA using the SE-TLD detector. All imaging was performed at 5mm working distance and in immersion mode. Horizontal field of view and dwell time were varied and are specified on each reported image. EDX measurements were performed in the same Nova Nanolab dual-beam SEM using an Oxford Instruments X-MAX 80 EDX detector. The working distance was kept at 5 mm to have the optimum EDX signal. Spectra were analyzed using the Oxford Instruments AZtec software. The reported map EDX was obtained with a primary beam energy of 5 kV. Si and Au are mapped and reported in weight percentage (Wt%).

### SEM image analysis

We aimed to isolate individual nanoparticles in SEM images to characterize them based on their core radius and tip length. Therefore, an algorithm was created using Python and OpenCV, available on GitHub.^51^ Firstly, the algorithm segmented the nanoparticles from the background using an Otsu adaptative threshold for bi-multimodal histograms. Secondly, an erosion was executed on the binary mask with 5 iterations and a 3 × 3 kernel to separate close nanoparticles. This way, we isolated the center of mass of the nanoparticles to iterate over the individual contours and generate individual masks for each nanoparticle; on the individual mask, we performed a symmetric dilatation to revert the previous erosion. Next, from the mask of the individual nanoparticles, we calculated the inner maximum radius to find the core size and the minimum enclosing circle to get the total radius and the tip length. Finally, we extracted the pixel/meters conversion factor using the images’ metadata for further analysis.

Aggregates that were still detected as single nanoparticles have been filtered post-analysis by applying the 2*σ* criterion on both radius and tip length.

### Fluorescence assays in solution

Experiments in **Figs. S9, 2f, 2g, 3** and **4** were performed with nanostars grown in-house; experiments in **Figs. 2b** and **2d** were performed with commercially available nanostars.

Commercial gold nanostars (100 nm) as well as biotin-functionalized gold nanostars (100 nm), were obtained from Cytodiagnostics (Catalog number: GU-100-100). Notably, their absorption maxima are closely aligned with our in-house synthesized nanostars. The experiments were conducted using standard 96-well plates. 10 nM nanostars or biotin-functionalized nanostars, Streptavidin-Cy5 (10 *µ*M, ThermoFisher Scientific) were introduced into the respective sample groups. The control group was prepared with six distinct replicates, and mixture and bound groups with three distinct replicates to ensure statistical robustness.

The 96-well plates, housing the nanostars and Streptavidin-Cy5 samples, were incubated at 37 °C for one hour to ensure complete binding between the components. Following this incubation, the fluorescence emissions were quantified and recorded with a Cary Eclipse Fluorescence Spectrophotometer (Agilent Technologies).

The statistical tests used for Fig. 2b are one-sided t-tests.

### Fluorescence spectroscopy assays on substrates

Glass coverslips underwent the MPTMS surface coating and nanoparticle immobilization, as indicated above. Approximately 60 *µ*L of this MPTMS solution was applied to the glass coverslips. The uniform layer of gold nanostars (AuNS) was deposited onto the substrates by adding 1 mL of AuNS solution to each recessed well.

The containers housing the glass slides were then sealed and placed on an orbital shaker (100 rpm) in a dark environment for an incubation period of 18 hours. On the following day, the AuNS-coated glass slides underwent three rinses with Milli-Q water, followed by gentle drying under a stream of nitrogen gas. To form monolayers of biotinylated bovine serum albumin (bBSA) on the substrates, 200 *µ*L of a 100 *µ*g/mL bBSA solution in PBS was applied to the substrate surfaces and allowed to incubate for 1 hour in a 37°C incubator. 40 *µ*L of a 10 *µ*M Streptavidin-Cy5 solution was added to the substrate surfaces and incubated for 2 hours in a 37 °C incubator. Following the incubation, the substrates were rinsed 3 times with PBS to eliminate any unbound fluorophores and gently dried using a nitrogen gas stream.

Fluorescence emission spectra of Streptavidin-Cy5 were measured using a photomultiplier tube (R7600U-20, Hamamatsu) coupled to a monochromator (SP2300, Princeton Instruments). Excitation light was provided by a xenon lamp connected to a monochromator (Gemini 180, HORIBA). The measurements were repeated only once for each group.

### Supported lipid bilayer formation

A glass slide covered by a silicone gasket with four 100 *µ*l wells was prepared with different MPTMS-immobilized nanostars and fibronectin conditions prior to the bilayer formation. The two Avanti Polar lipids (Birmingham, AL) used were 1,2-dioleoyl-sn-glycero-3-phosphocholine (DOPC) and 1,2-dioleoyl-sn-glycero-3-phosphocholine-N-(Cyanine 5) (DOPC-Cy5). Lipids in chloroform were mixed in a molar ratio of 99.5:0.5 DOPC:DOPC-Cy5 in a glass vial. The chloroform was evaporated with nitrogen and dried in a vacuum desiccator overnight. The dried lipid film was resuspended in PBS. The glass vial with the solution was then vortexed in bursts of 30 seconds with 1-minute intervals to resuspend all lipids and to stimulate the self-assembly of multilamellar vesicles. The solution was then sonicated in a high-capacity bath sonicator (10% amplitude, 10 s on, 10 s off, for one hour) to get a uniform size distribution of liposomes. 100 *µ*l of liposome solution was then deposited in each of the wells of the silicone gasket. The glass slides with liposomes were incubated for 30 min at room temperature to allow for supported lipid bilayer formation. All wells were washed four times with 200ul PBS after incubation.

### TIRF microscopy

For the TIRF microscopy, an inverted microscope (NikonTi2-E, motorized) equipped with a 100× oil immersion objective (Nikon Apo TIRF 1.49 NA) and an Andor iXon Ultra 897 camera was used. The microscope was upgraded with an azimuthal TIRF/FRAP illumination module (GATACA systems iLAS 2). Samples were exited at 642 nm at 10% laser power (GATACA CW laser, 642nm/110mW). The emission was filtered with a 706/90 filter. Samples were exited for 200 ms for each frame, and a multiplication gain 50 was used. The penetration depth automatically estimated by the system is 106 nm. Every group consists of 21 distinct samples.

### HEK293T culture

HEK293T cells (Catalog number ATCC® CRL-3216™) with passage number ¡30 were cultured in Dulbecco’s Modified Eagle Medium (DMEM) supplemented with 10% Fetal Bovine Serum (VWR Seradigm Premium Grade), 50 units/mL Penicillin (Life Technologies), 50 *µ*g/mL Streptomycin (Life Technologies), and 1% GlutaMAX (Life Technologies) at 37 °C under 10% CO_2_.

### Lentivirus preparation and transduction

For lentivirus production, HEK293T cells were seeded at 4*·*10^6^ cells per 100 mm polystyrene dish (Sarstedt) 1 day before transfection in order to achieve 80% confluency at the moment of transfection. Cells were co-transfected with 1.5 *µ*g transfer plasmid carrying either QuasAr1 or QuasAr6a, 5 *µ*g of pMDLg/RRE (Addgene plasmid #12251), 5 *µ*g of pRSV/Rev (Addgene plasmid #12253), and 3.5 *µ*g of pMD2.G (Addgene plasmid #12259) mixed in 400 *µ*L pre-warmed opti-MEM with 60 *µ*L polyethylenimine solution (1 *µ*g/*µ*L in PBS). 8 hours post-transfection, medium was replaced with fresh cell culture medium including 25 mM HEPES. Lentivirus-containing supernatant was collected 48 and 72 hours post-transfection. Supernatant collected 48 hours post-transfection was stored overnight at 4 °C before pooling it with lentivirus collected 72 hours post-transfection. Supernatant was filtered through 0.22 *µ*m PVDF filters. Lentivirus was concentrated 15-fold using LentiX concentrator reagent (Takara), resuspended in PBS, and stored at -80 °C in 100 *µ*L aliquots. *µ* For lentiviral transduction, 2.5*·*10^5^ cells were seeded per 35 mm polystyrene dish (Sarstedt) 1 day before transduction in order to achieve 50% confluency at the moment of transduction. Lentiviral aliquots were rapidly thawed in a 37 °C water bath. 100 *µ*L of concentrated virus was added per dish. Cells were incubated with lentivirus for 24 hours, after which the media was replaced with fresh cell culture medium.

### Widefield fluorescence microscopy

A custom-built^52^ widefield microscope was employed to investigate the fluorescence of the QuasArs expressed in cells. Two continuous wave laser sources (MLL-FN-639 CNI and OBIS 488 LX, by Coherent) were made uniform in polarization and beam diameter through zero-order half-wave plates (WPH05M-488 and WPH05M-633 by Thorlabs), polarizers (CCM5-PBS201/M by Thorlabs) and individual collimators (AC254 mounted achromatic doublets by Thorlabs). The beams were then combined using a dichroic mirror (DMLP505 by Thorlabs) and led through an acousto-optic tunable filter (AOTFnC-VI S by AA Optoelectronics) with a modulation rate of 22 kHz. With the exception of the patch clamp voltage imaging experiments, the beam diameter was expanded with a telescope (AC254 mounted achromatic doublets by Thorlabs) using flip mounts (TRF90/M by Thorlabs). The light was reflected from a Digital Micromirror Device (DMD, Vialux V7001 by Texas Instruments) into a tube lens (TTL200 by Thorlabs) and projected by the objective (Olympus XLPLN25XWMP2, NA 1.05, working distance 2 mm) onto the sample. The emitted fluorescence was reflected by a dichroic mirror (Di03-R405/488/532/635-t3-32 × 44 by Semrock), collected using a tube lens (TTL200 by Thorlabs) and filtered further depending on the emission range (FF01560/94-25, FF01-582/64-25, LP02-664RU-25 and FF01-790/SP-25 by Semrock), before reaching the sCMOS camera (ORCA Flash4.0 V3, 2048 × 2048 pixels, 6.5 *µ*m pixel size by Hamamatsu) where the image was captured in a low-noise configuration with a total magnification of 27.8. The maximum powers after the objective in the telescope configuration are 40 mW for the 639 nm laser and 2 mW for the 488 nm laser; without the telescope, the powers are 122 mW for the 639 nm laser and 5.2 mW for the 488 nm laser.

In Fig. 3a the sample size is n=4 independent cells for the control and n=4 independent cells for the nanostar sample. In Fig. 3b the sample size is n=10 independent cells for the control and n=8 independent cells for the nanostar sample. In Fig. 3e the sample size is n=73 independent cells for the control and n=180 independent cells for the nanostar sample.

The statistical test used for Fig. 3e is a one-sided Mann-Whitney test that does not assume a Gaussian distribution of the data points.

### Whole-cell patch-clamp recordings

Patch-clamp and imaging were performed 72 hours after lentiviral transduction. Transduced cells were seeded in the 14mm glass well of fibronectin-coated glass bottom dishes (Cellvis) at a density of 1.5*·*10^4^ cells per well 24 hours before the patch-clamp experiment. Before patchclamp procedure, cells were washed gently three times with 2 mL of pre-warmed extracellular buffer (EC buffer). Cells were immersed in 2 mL of EC buffer throughout the patch clamp procedure. Constitution of EC buffer is 125 mM NaCl, 2.5 mM KCl, 3 mM CaCl2, 1 mM MgCl2, 15 mM HEPES, and 30 mM glucose.^53^ pH of EC buffer is adjusted to 7.3 with NaOH. Osmolarity of EC buffer is adjusted to 300 mOsm with sucrose.

Borosilicate glass micropipettes were pulled from capillaries (Sutter Instrument) and filled with 10 *µ*L of intracellular medium (IC medium). Constitution of IC medium is 125 mM K-gluconate, 8 mM NaCl, 0.6 mM MgCl2, 0.1 mM CaCl2, 1 mM EGTA, 10 mM HEPES, 4 mM Mg-ATP and 0.4 mM Na-GTP, with pH adjusted to 7.3 with KOH.^54^ Osmolarity of IC medium is adjusted to 295 mOsm with sucrose. Tip resistance was verified to be between 5 and 10 MΩ. Cellular responses were evoked through voltage steps ranging from -70 mV to +30 mV at a frequency of 5 Hz, while the fluorescence of QuasAr1 and QuasAr6a was excited at 639 nm. The camera frame rate in this setting is 1000 fps.

In Fig. 4 the sample size is n=7 independent cells for the control and n=5 independent cells for the nanostar sample. The statistical tests used for Fig. 4c,d are one-sided t-tests.

The photocycle of QuasAr6a has been modelled as a 4-state continuous-time Markov chain, where the putative biological macrostates corresponded directly to the mathematical states. Transitions were considered unidirectional. The Markov chain was expressed as a system of linear ODEs and solved numerically in Python using the LSODA solver (Livermore Solver for Ordinary Differential equations with Automatic method switching for stiff and non-stiff problems). The features of simulated traces (brightness enhancement, voltage sensitivities, response speeds) are compared to those of the measured traces and improved following a differential evolution algorithm. The software outputs the best parameters after a maximum of 1000 iterations, unless a plateau is reached earlier. To quantify the goodness of the fit in different conditions, we employed a Huber loss calculated on the fitted features compared to the one from the simulations. A lower loss indicates a better agreement between the simulations and the data. The fitted values for the models can be consulted in the Supplementary Information, while the code can be accessed on GitHub.^51^

### AI Methods

Various AI-based tools were used to aid in the creation of this manuscript.

ChatGPT (GPT-3.5 and GPT-4o) was used to speed up and optimize the coding process in Python. All AI-generated code was debugged and validated.

Paperpal, Typeset.io and Research Rabbit were used to assist in finding papers relevant to the current manuscript for the literature review. ChatGPT was used to check consistency between the manuscript text and cited references. As with any other library-building tool, all suggested papers were reviewed manually regarding their relevance and accuracy prior to the decision to cite a reference in the manuscript.

ChatGPT was used to check flow of paragraphs and phrasings of specific sentences in a first draft of this manuscript, before complete revision by authors. Grammarly was used for grammar checks, correcting errors and enhancing clarity of the text. Suggestions were considered critically, especially if changes in punctuation or syntax could distort the original message conveyed in the text.

Any intellectual contributions, final decisions on content and structuring of the work, and writing of the final manuscript were exclusively made and done by credited authors.

## Authors’ contributions

**Table 1:**
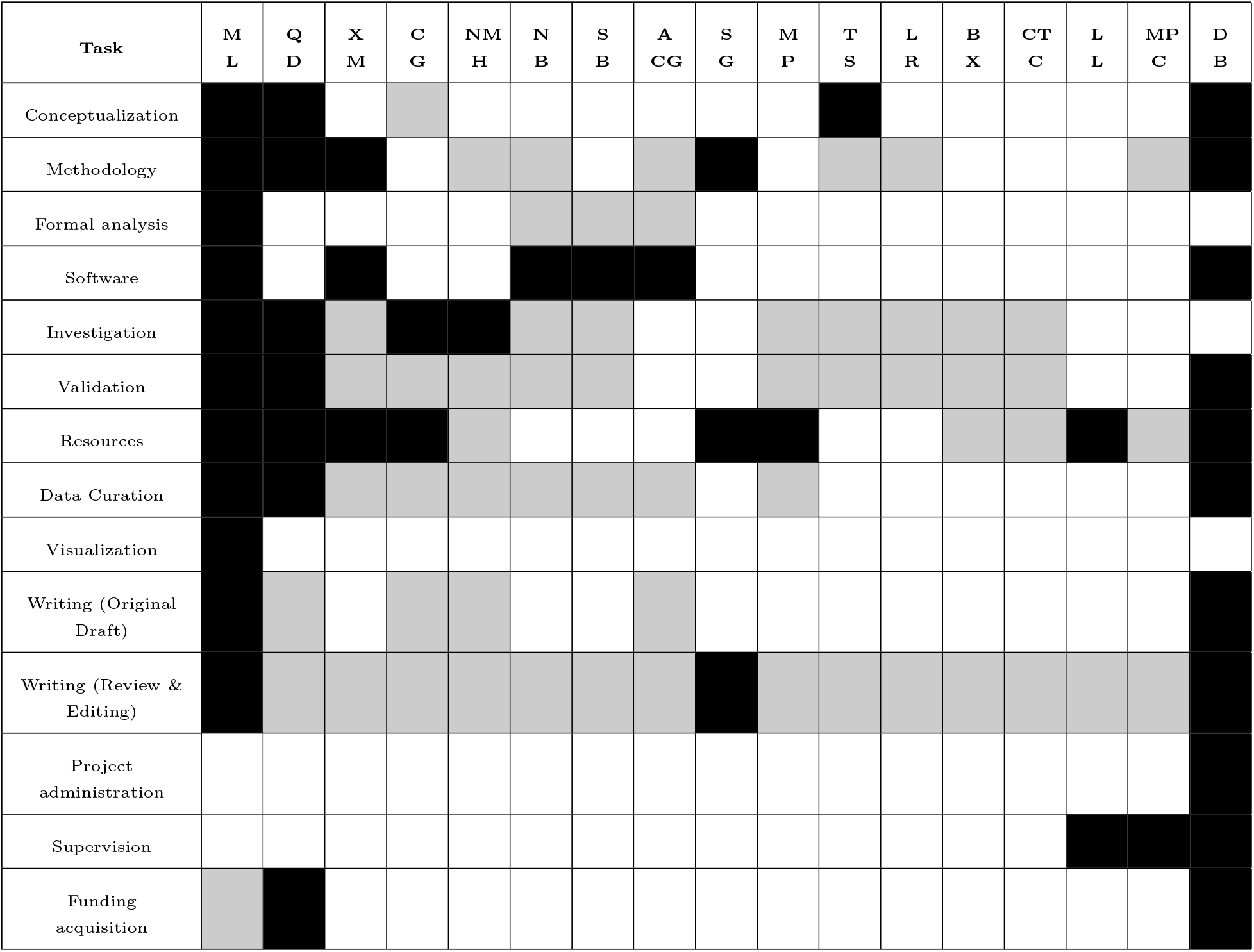
Authors’ contributions according to CRediT taxonomy.

## Acknowledgments

We thank Ryan Lane for preliminary SEM images that did not make it to the final draft, Zimu Wei for UV-Visible spectroscopy measurements that did not make it to the final draft, Zhenzhen Wu for lifetime experiments that did not make it to the final draft, Ornella Cavalleri and Claudio Canale for suggesting the use of MPTMS, Antonia Denkova and Erik van der Kolk for training and use of facilities. We thank Adam Cohen and Tian He for generously sharing QuasAr1 and QuasAr6a with us. ML acknowledges support from the Erasmus+ Traineeship programme. QD acknowledges support from the Chinese Scholarship Council scholarship. DB acknowledges support from an NWO Start-up Grant (740.018.018) and ERC Starting Grant (850818 - MULTI-VIsion), as well as an NWO XS grant (OCENW.XS2.033), and by the Delft AI initiative (Ailab BIOlab) and the Convergence Health and Technology (Integrative Neuromedicine flagship).

## Ethics and inclusion statement

The research in this paper does not involve local researchers. All biological material was obtained according to the Nagoya protocol.

## Data availability statement

All data underlying manuscript figures have been uploaded to the 4TU Respository and are available at DOI: 10.4121/909b7170-816f-4d81-a8b0-12b113f29207

## Code availability statement

The code running the setup on which these measurements were done is available at: https://github.com/Brinkslab/gevidaq.

The code used for the analysis of the data is available at: https://github.com/Brinkslab/plasmon_gevi.

