## Supplemental information for "Plasmonic Enhancement of Protein Function"

### Supplementary Information:
### Plasmonic Enhancement of Protein Function

Marco Locarno,† Qiangrui Dong,† Xin Meng,† Cristiano Glessi,† Nynke Marije Hettema,‡ Nidas Brandsma,† Sebbe Blokhuizen,† Alejandro Castañeda Garcia,† Srividya Ganapathy,†,¶ Marco Post,† Thieme Schmidt,† Lars van Roemburg,† Bing Xu,§ Chun-Ting Cho,§ Liedewij Laan,‡ Miao-Ping Chien,‖ and Daan Brinks*,†,‖

†*Department of Imaging Physics, Delft University of Technology, Delft, The Netherlands*

‡*Department of Bionanoscience, Delft University of Technology, Delft, The Netherlands*

¶*Department of Pediatrics and Cellular and Molecular Medicine, UCSD School of Medicine, San Diego, California, United States*

§*Department of Radiation Science and Technology, Delft University of Technology, Delft, The Netherlands*

‖*Department of Molecular Genetics, Erasmus University Medical Center, Rotterdam, The Netherlands*

### FDTD simulations

#### Spherical gold nanoparticles

Numerical simulations of gold nanospheres were conducted using the script for Lumerical's FDTD Solutions, which is publicly accessible in the GitHub repository.[1]

The code performs simulations by varying the radius $R$ of a sphere submerged in water. It identifies the maximum field enhancement $|E|^2/|E^0|^2$, averaged within a small volume ($\Delta x = R/2$, $\Delta y = R/2$, $\Delta z = R/2$) located 4 nm away from the surface. The resonance wavelength and the corresponding average field enhancement are then recorded in a text file. This computational "screening" process continues within a specified range, with steps of 2 nm, ranging from $R_{\mathrm{min}} = 6$ nm to $R_{\mathrm{max}} = 76$ nm.

**Figure S1** illustrates the relationship between the sphere's radius and the peak wavelength in the average field enhancement. The simulation results align well with the theoretical model of the Modified Long-Wavelength Approximation (MLWA),[2] indicating the limited tunability of the plasmonic resonance for this simple morphology. This red-shift can be attributed to dynamic depolarization of the nanoparticle.[2]

The dampening and broadening of the field enhancement for particles with radii larger than 30 nm can be attributed to the radiative damping arising from the spontaneous emission of radiation by the induced dipole.[2] These simulations are valuable for verifying the reliability of the employed method and directly computing the local electric field instead of relying on indirect estimates.

The initial findings have brought to light the importance of finding a shape that offers a more tunable and intense plasmonic resonance. This realization has led us to explore the potential of utilizing the peak splitting observed in ellipsoids. The idea is then to identify a shape similar to the ellipsoid, but also practical and straightforward to fabricate with techniques like electron beam lithography.

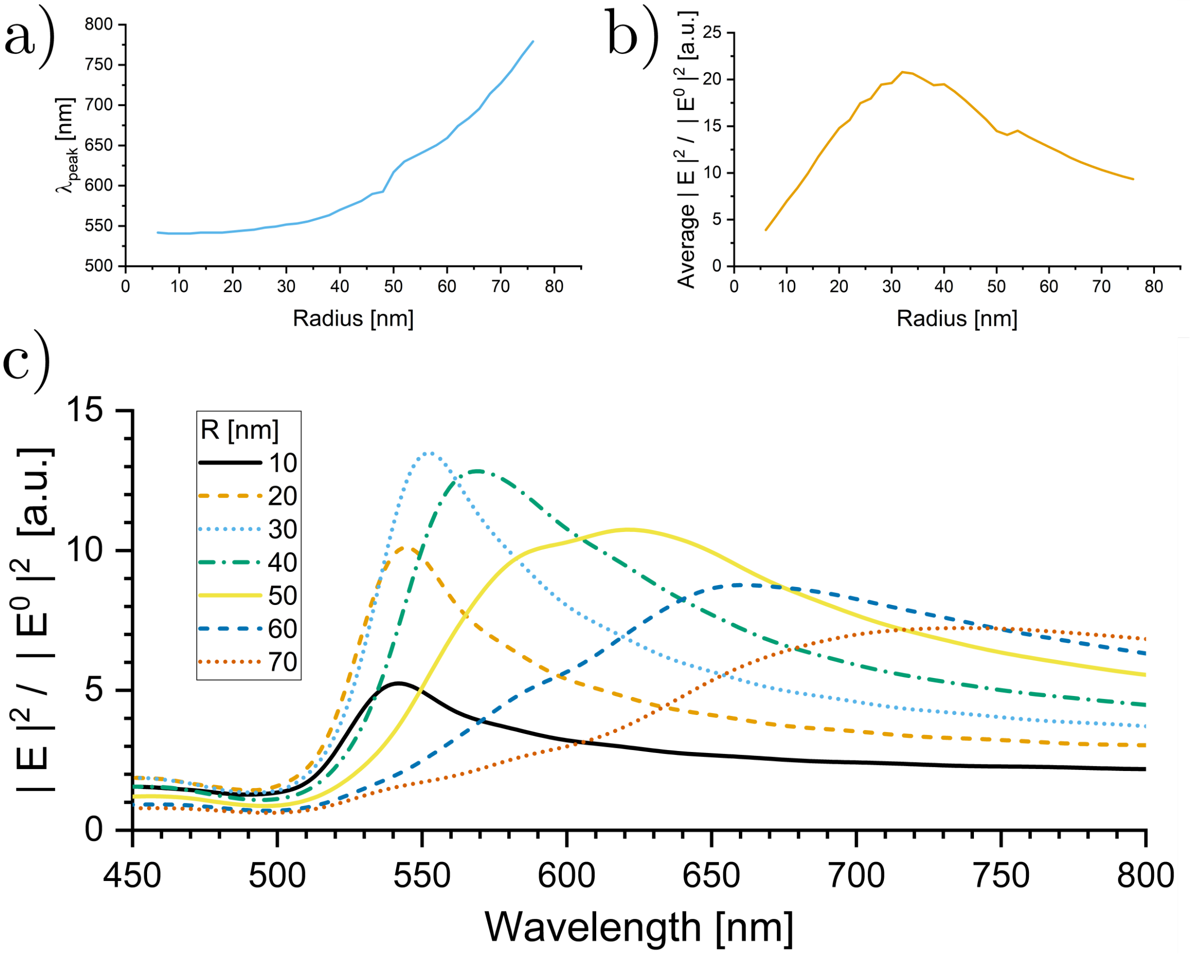


**Figure S1: a,** Peak position and **b,** intensity of the average field enhancement of gold nanospheres in water, measured in a volume 4 nm away from the surface of the particle, as a function of radius. Larger radii correspond to stronger peak's redshift and lower field enhancement. **c,** Spectra of gold nanospheres immersed in water for different radii. The smaller nanoparticles (10-40 nm) have a resonance peak around 550 nm. The resonance peaks are redshifted for the larger particles, with some even going into the IR region. For particles larger than $\sim 30$ nm, the width of the peak is significantly broadened.

#### Smooth gold nanorods

In this section, we focus on realistic nanorods that do not have very sharp edges. In a sense, they are similar to ellipsoids, but present flatter surfaces and have overall larger volumes. These nanorods are particularly interesting due to their unique plasmonic properties and tunability, and could be nanofabricated in regular patterns.

Numerical simulations of gold smoothed nanorods were performed using the script for

Lumerical's FDTD Solutions, which is publicly available in the GitHub repository.[1]

In brief, the code varies the three main parameters (namely length $l$, width $w$ and height $H$) of the smoothed rod immersed in water and computes the maximum average field enhancement $|E|^2/|E^0|^2$ in a small volume ($\Delta x = w/2$,$\Delta y = w/2$,$\Delta z = H$) 4 nm away from the surface. Then, the resonance wavelength and the corresponding average field enhancement are saved in a text file. This computational screening continues in a defined range by steps of 5 nm:

- 20 nm $\leq l \leq$ 100 nm;
- 20 nm $\leq w \leq$ 50 nm;
- 20 nm $\leq H \leq$ 50 nm;

To save computational time, the script skips tests with higher $l$ if the resonant wavelength is longer than 780 nm (outside the region of interest).

In **Fig. S2**, each data point corresponds to the field enhancement of an individual nanorod with a fixed combination of dimensions ($l$, $w$, $H$). The ranges chosen for the simulations focused solely on the longitudinal resonance, resulting in the absence of the transverse resonance in the graph. As anticipated by the theory of elongated particles[2] and consistent with previous nanorod experiments,[3] the field enhancement becomes stronger as the resonance is red-shifted. This behaviour is directly related to the aspect ratio of the nanoparticle.

Notably, these simulations deviate from other studies by taking into account various values for the height $H$ rather than treating it as negligible. The effect of height is qualitatively similar to that of changing the width, contributing to multiple combinations of parameters yielding similar resonance wavelengths.

While the smooth gold nanorods exhibit tunability within a desirable range (near-IR), two main challenges arise with this morphology. First, when these nanorods are nanofabricated on a substrate, the field enhancement remains confined to that plane. Given that biological

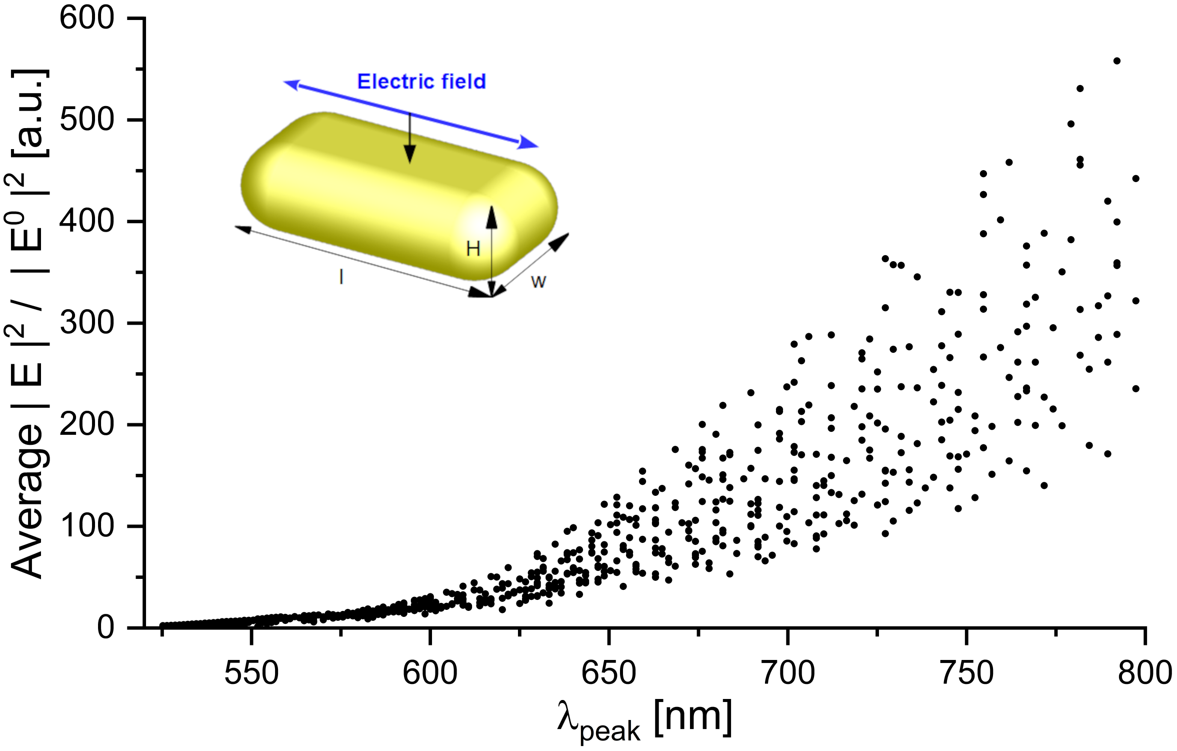


**Figure S2:** Results of the computational screening of gold nanorods, where each spot corresponds to a set ($l$,$w$,$H$).

applications occur in complex three-dimensional environments, this limitation poses a significant obstacle. On the other hand, if the nanorods are synthesized in a colloidal solution, achieving precise control over their orientation becomes difficult. Consequently, there is a risk of exciting the blue-shifted transverse mode instead of the more intense longitudinal mode.

To address these limitations and build upon the concept of elongated features, we look to spiked nanoparticles as the next logical step. By combining the elongated aspect with the rotational symmetry of spheres, spiked nanoparticles offer promising possibilities. These structures hold the potential to overcome the challenges faced by smooth nanorods and provide enhanced control over plasmonic resonances in three-dimensional environments.

#### Octahedral silver nanoparticles

Having established a solid understanding of plasmonic resonance in smooth nanoparticles, we can now explore the realm of nanostars, which offer even more significant local enhancement thanks to their sharp tips. An octahedral nanostar structure was employed to address the issue of rotational dependence in the enhancement. This particle exhibits rotational symmetry in multiple directions, simplifying the simulation process. Instead of varying multiple rotation angles, only one angle needed to be adjusted to capture the complete 360-degree behaviour of the enhancement.

For this section, the material of the nanoparticles was arbitrarily chosen as silver instead of gold, thanks to the input from the supervised student who contributed to this research. Since our focus is primarily on the impact of shape, the choice of material should have minimal influence on our analysis and treatment.

Numerical simulations of silver octahedral nanostars were performed using the script for Lumerical's FDTD Solutions, which is publicly available in the GitHub repository.[1] The conical base was set to have a conical angle of either 25 degrees or 40 degrees. To create the rounded tips, the truncated cone was connected to a semisphere, with the semisphere's radius set to 0.05 times the particle's radius ($0.05R$). Additionally, the length of the spikes was determined to be 1.5 times the particle's radius ($1.5R$). This conical structure more realistically represents a physical nanostar compared to a cone with an infinitely sharp tip.

In order to assess the bandwidth of the enhancement, simulations were conducted by rotating the particle between 0 and 45 degrees. Further rotations beyond this range were deemed redundant due to the 4-fold symmetry of the particle. Throughout the simulations, the monitor remained fixed to maintain consistency. This comparative analysis aimed to provide insights into the distinctive properties and performance of the nanostar structure as a function of rotation.

As visible in **Fig. S3a**, there is a strong dependence of the field enhancement intensity on the rotation of the particle. However, the wavelength at which the plasmonic resonance

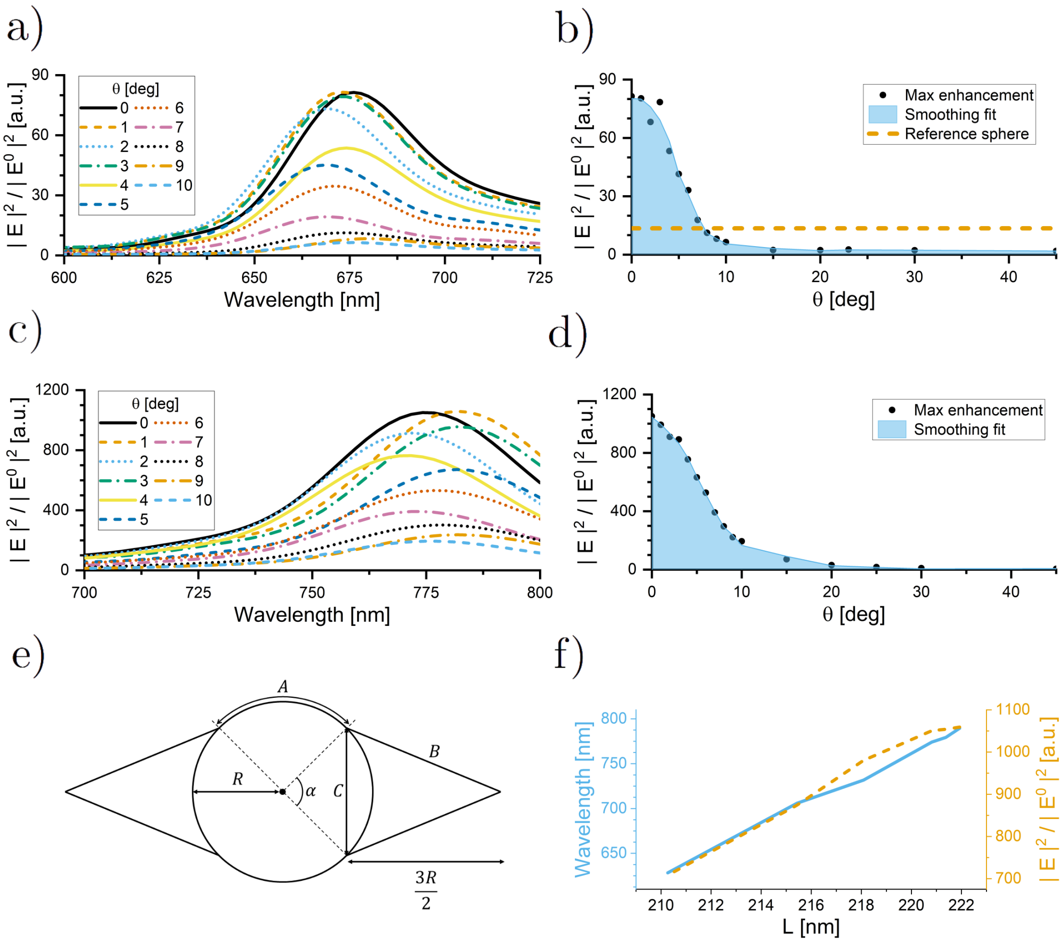


**Figure S3:** Angles don't significatively change the peak position of the plasmonic resonance, but they affect the local field on-axis. Larger base angles result in red-shifted and more intense plasmonic peaks. Both peak position and field enhancement are linearly dependent on the circumference in the observed plane.

occurs is very stable. Here, the centre spherical particle had a radius of 20 nm, and the spikes had a length of 30 nm; therefore, the characteristic length of the nanostar is $2 \cdot (20+30) = 100$ nm. Therefore, we compare it to a spherical nanoparticle with a diameter of 100 nm to get a rough comparison for the practicality of the nanostar. We want the average enhancement of the nanostar to be higher than the spherical silver nanoparticle.

By calculating the average enhancement per degree over the entire range of rotation, we find an average enhancement of 12 per degree. Since the enhancement of the spherical particle does not change under rotation, the average enhancement is simply the maximum enhancement, so 13.5 per degree. We observe that, on average, the enhancement of the nanostar is less high than the spherical particle in the visible spectrum. In an effort to increase this bandwidth, we tested a different conical base angle of the spikes whilst keeping the radius and spike length the same (**Fig. S3c**).

The results reveal a remarkable enhancement peak in the nanostar structure, surpassing previous simulation findings by over an order of magnitude. Additionally, similar to the earlier observations, the wavelength corresponding to this higher enhancement peak remains relatively stable even when the particle is rotated. Regrettably, the spectrum bandwidth remains essentially unchanged compared to the previous simulations. However, the significant increase in enhancement values leads to a noteworthy rise in average enhancement per degree, reaching an impressive value of 167 per degree. Compared to its spherical counterpart with a similar characteristic length (as discussed earlier), the nanostar exhibits a substantially higher average enhancement.

The characteristic length so calculated is independent of the conical aperture of the spikes, therefore not explaining the different behaviours in **Figs. S3a** and **S3c**. A different way to evaluate the characteristic length is calculated according to the definitions in **Fig. S3e** as the circumference (in the section plane) of the particle. The perpendicular spikes are neglected as they do not affect the plasmonic enhancement (transverse mode).

The formula used to calculate the characteristic length, with $\alpha$ in radians, is:

$$L = 2R\left[\pi - \alpha + 2\sqrt{\sin^2\left(\frac{\alpha}{2}\right) + \frac{9}{4}}\right] \tag{S.1}$$

We considered conical base angles ranging from 38 to 60 degrees, as these angles correspond to peaks within the visible spectrum. Using **Eq. (S.1)**, we calculate the circumference lengths and plot them against both the resonance wavelength and the maximum enhancement in **Fig. S3f**. Both the maximum enhancement and the peak resonance wavelength exhibit a linear relationship with the circumference length.

**Figure S4** clearly demonstrates the persistence of an intense plasmonic effect in the spiked nanoparticle; however, the localization of this effect at the tip implies that the measured field within a fixed window will significantly decrease as the nanoparticle undergoes rotation.

Given the orientation-dependent nature of octahedral nanoparticles, we contemplated the idea of utilizing a spiked nanoparticle with a high density of tips as a potential solution to overcome this challenge. The abundant spikes in such a structure could offer advantages in terms of orientation insensitivity, allowing us to explore plasmonic properties more effectively.

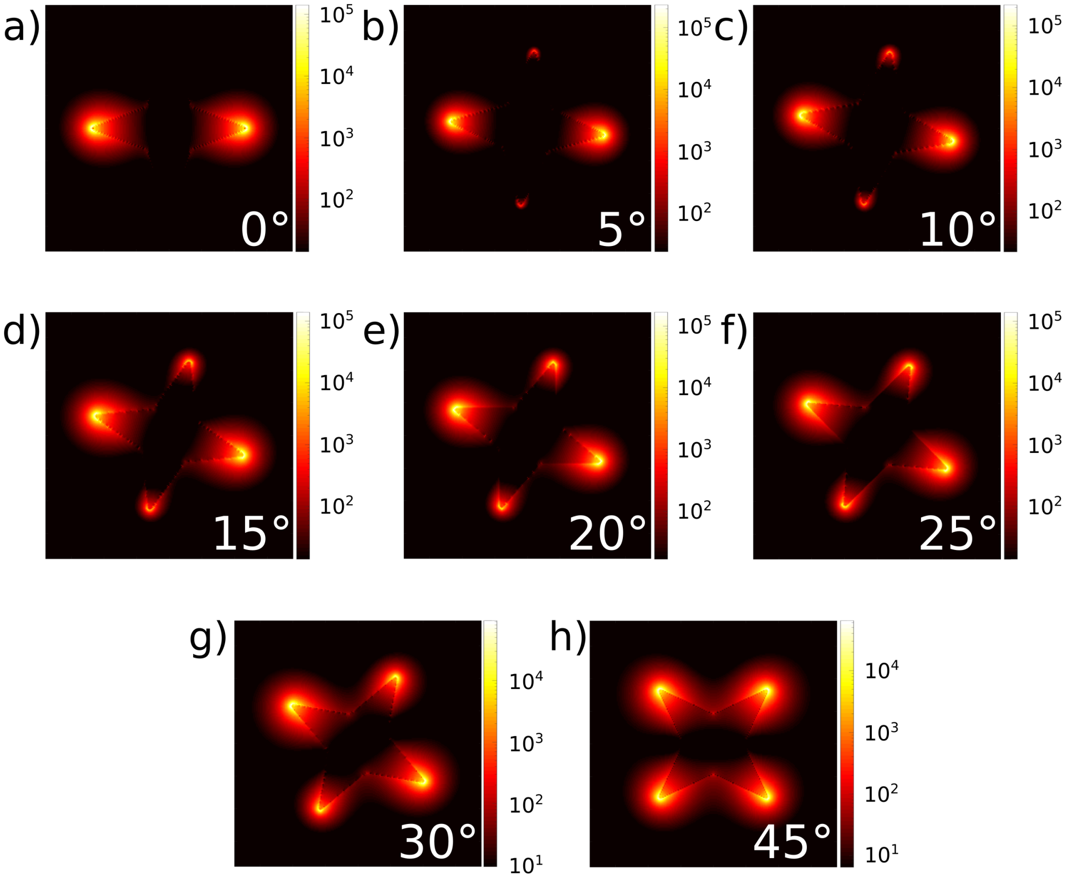


**Figure S4:** Field enhancement maps for a silver octahedral nanostar with 40 degrees of conical angle, for different rotations, at the resonance wavelength (774 nm). The maximum enhancement close to the tip is roughly the same order of magnitude, but this localization results in a lower field enhancement on the x-axis for some rotation angles.

#### Highly spiked gold nanostars

Numerical simulations of highly spiked gold nanostars were conducted using the script for Lumerical's FDTD Solutions, which is publicly available in the GitHub repository.[1]

In brief, the code performs simulations by varying the core radius $R$ of the nanostar immersed in water, keeping the tip length fixed at 10 nm. The primary goal is to compute the average field enhancement $|E|^2/|E^0|^2$ within a small volume ($\Delta x = R_{\mathrm{core}}$,$\Delta y = R_{\mathrm{core}}$,$\Delta z = R_{\mathrm{core}}$) positioned 4 nm away from the surface. Subsequently, the resonance wavelength and the corresponding average field enhancement values are recorded in a text file. Additionally, the code saves the absorption and scattering cross-sections as functions of the wavelength. The simulations cover a defined range, incrementing the core radius by 5 nm, spanning from a minimum core radius of $R_{\mathrm{core,min}} = 20$ nm to a maximum core radius of $R_{\mathrm{core,max}} = 50$ nm.

The presence of the tips in the nanostars, as anticipated from the theory,[4] leads to an immediate red shift of approximately 100 nm compared to the plasmonic resonance observed in gold spherical particles of the same size. Unlike nanospheres, nanostars exhibit a more limited shift in resonance wavelength when the core size is altered, allowing for finer tunability of the peak. The initial blue shift is attributed to the competing effects of absorption and scattering, as illustrated in **Fig. S5**.

Interestingly, due to the presence of the spikes, the absorption peak consistently appears blueshifted relative to the scattering peak. Consequently, this results in a general blue shift when larger particles are grown, followed by a subsequent red shift. **Figure S5** offers a comprehensive comparison, outlining the primary effects of growing larger nanostars. The most notable effect is the significant broadening of the peak, resulting from non-dipolar effects, which leads to a less selective plasmonic resonance.

The simulation work focusing on highly spiked nanostars, characterized by a high density of tips around the spherical core, primarily remained theoretical. Nevertheless, these simulations already indicated the potential of such a morphology in achieving a favourable combination of resonance control, field enhancement engineering, and insensitivity to rota-

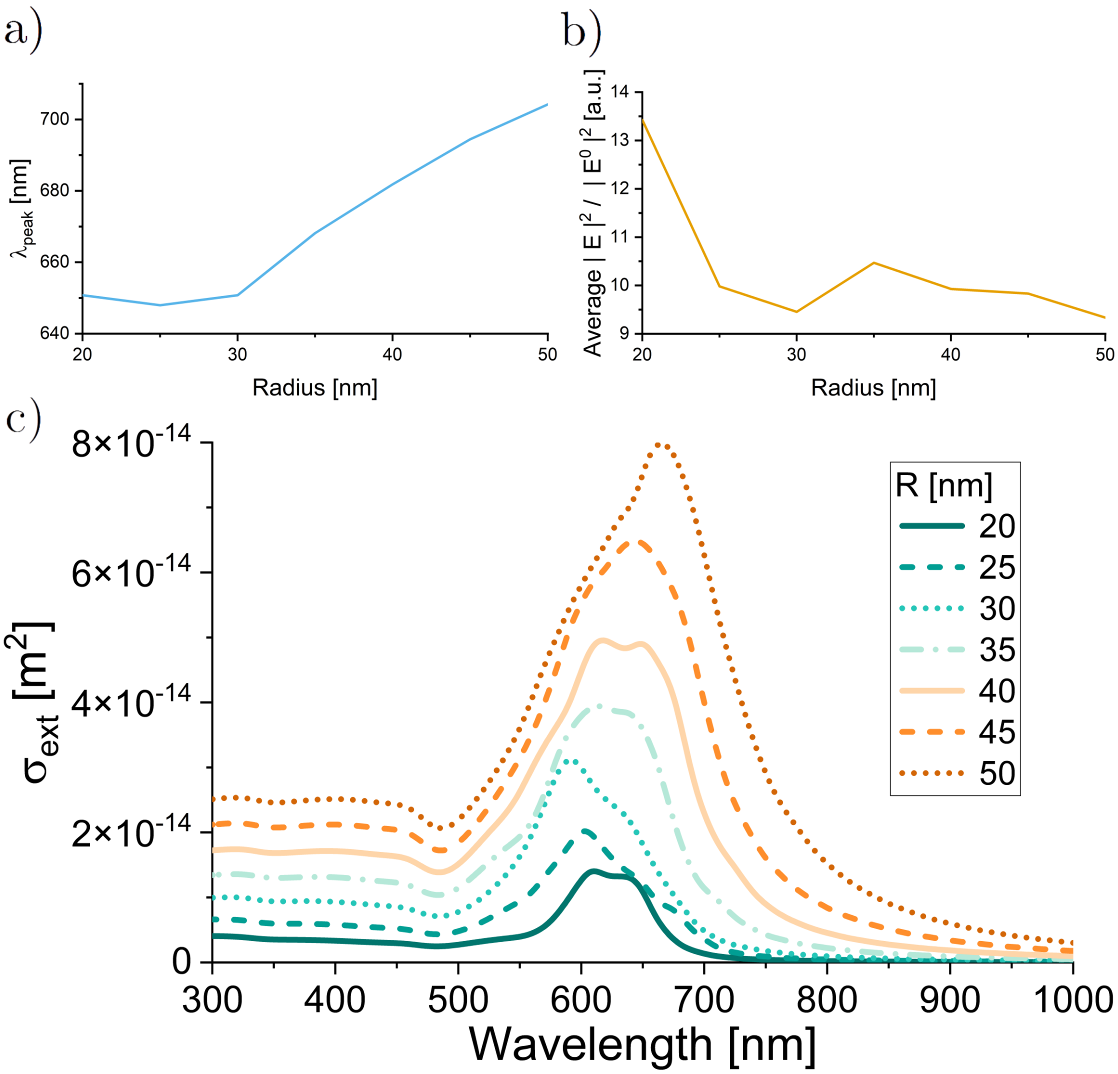


**Figure S5:** Screening of gold nanostars (high spike density) in water; larger radii correspond to stronger peak's redshift. Field enhancement is not significantly dependent on core size.

tions. To make the study more relevant and applicable to real-world scenarios, it became essential to identify an existing synthesis protocol capable of producing gold nanostars with the plasmonic peak positioned in the red region of the visible range. Thus, to bridge the gap between theory and practicality, we adapted and applied these simulations to realistic gold nanostars as discussed in the main manuscript.

### Individual simulations realistic nanostars

Here reported are the individual simulation results, which were then aggregated in **Fig. 1d** and **1e** (main manuscript).

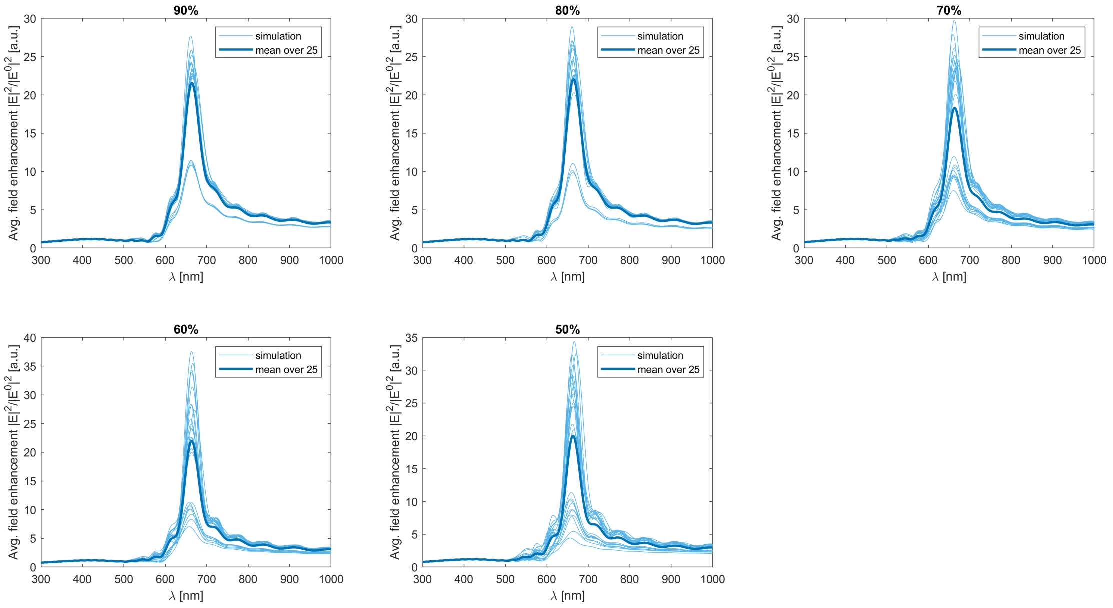


**Figure S6:** Invididual simulation results at different tip lengths. The results are aggregated in **Fig. 1d**.

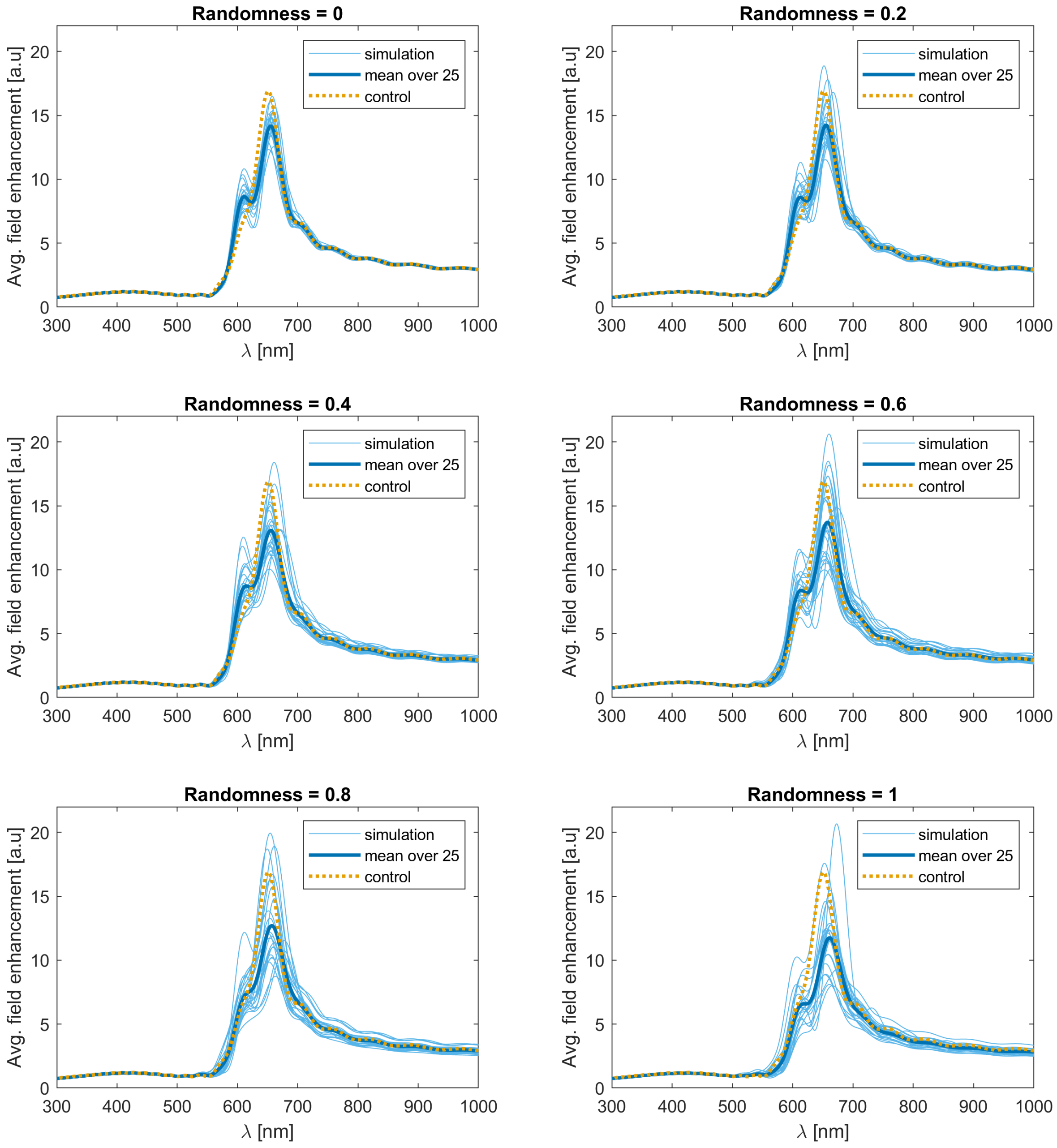


**Figure S7:** Invididual simulation results at different randomness factors. The results are aggregated in **Fig. 1e**. The control corresponds to an ideal nanostar, optimally aligned with the field monitor.

#### Energy-dispersive X-ray spectroscopy

Energy-dispersive X-ray (EDX) spectroscopy performed on the synthesized samples substantiated the purity of the gold nanostars, with no residual gold detected in the sample (**Fig. S8**).

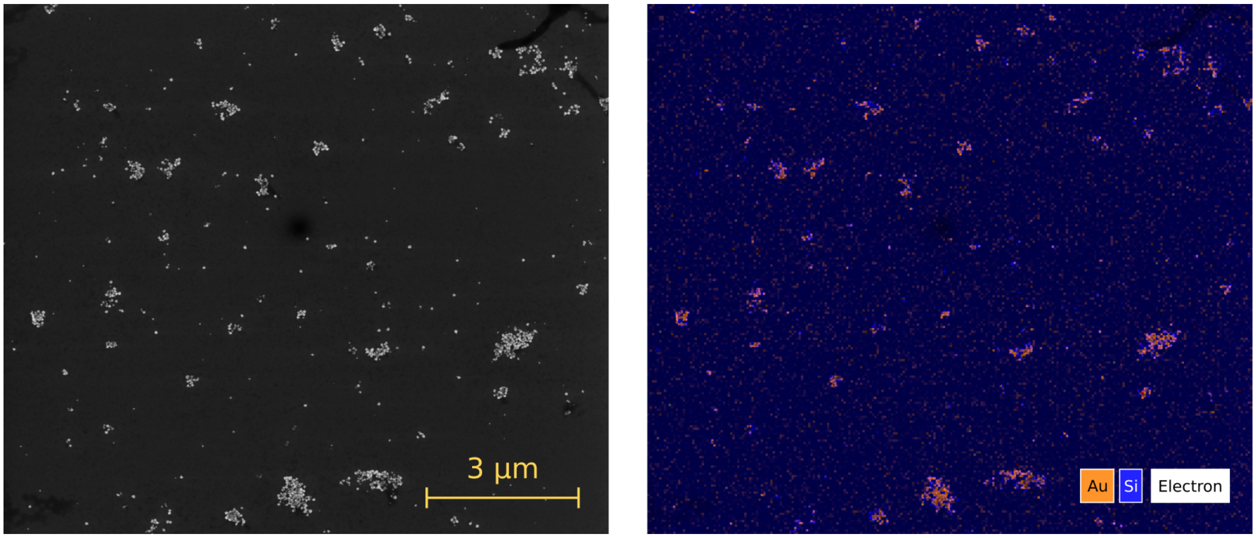


**Figure S8:** Validation of the purity of gold nanostar samples, showing the absence of any residual gold in the specimen. On the left, the SEM micrograph of a nanostar sample bound to glass via MPTMS; on the right, the same field of view under EDX inspection.

### Colloidally grown nanostars

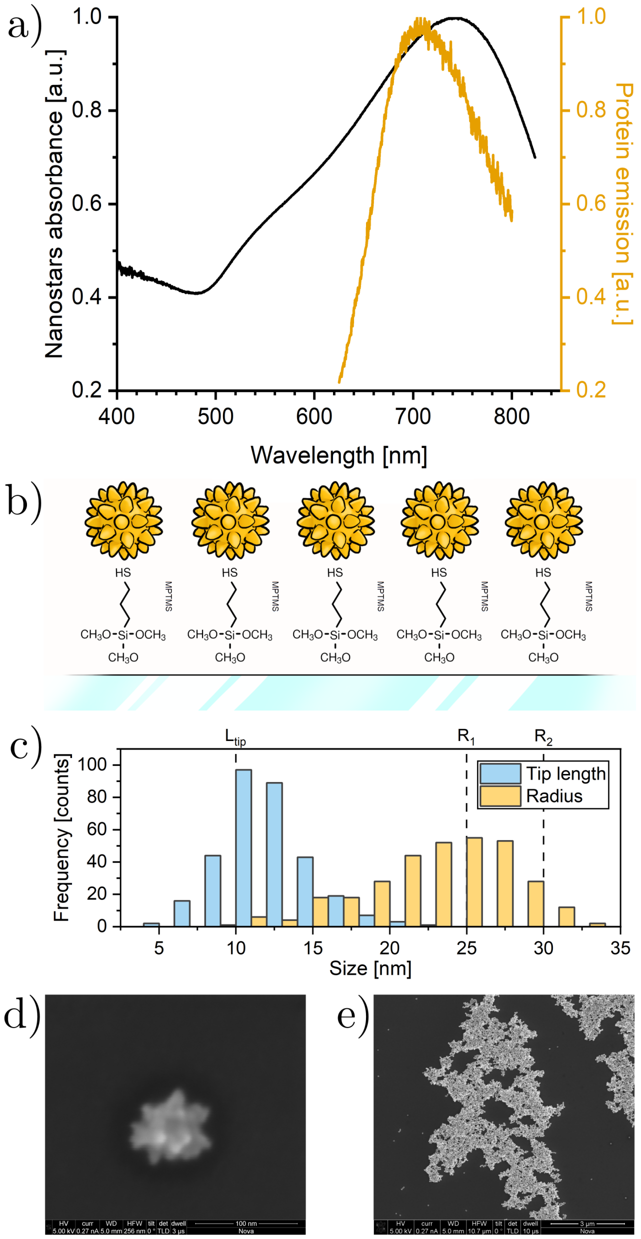


**Figure S9:** Colloidal gold nanostars are designed to enhance the fluorescence of QuasArs. **a,** Spectral alignment between the normalized absorbance of the gold nanostars colloid and the fluorescence emission of Archon1.[5] **b,** Illustration of the gold nanostars bound to glass via MPTMS. Not in scale. **c,** The physical features of the nanoparticles in SEM images match the ones used for the simulations in **Fig. 1**. $L_{tip} = 10$ nm is the tip length used in the simulation, and $R_1 = 25$ nm and $R_2 = 30$ nm are the two radii used in different simulations. **d,** SEM micrograph of an isolated gold nanostar. **e,** SEM micrograph of a large, complex cluster of gold nanostars.

#### Fröhlich-like shift due to PVP coating

The local dielectric environment has an effect on the plasmonic resonance frequency. To estimate this in the case of PVP-capped nanoparticles, let us assume that the red-shift due to the capping agent is Fröhlich-like:[2]

$$\lambda_{PVP} = \frac{2\pi c\sqrt{\epsilon_{m,\infty} + 2\epsilon_{PVP}}}{\omega_P} \tag{S.2}$$

where $\lambda_{PVP}$ is the resonance wavelength when considering the PVP as a capping agent, $c$ the speed of light, $\epsilon_{m,\infty}$ the core polarization of gold, $\epsilon_{PVP}$ the dielectric constant of PVP and $\omega_P$ the plasma frequency of gold.

If the nanostars would be "bare", or simply said immersed in water without any capping agent, the resonance wavelength would be:

$$\lambda_{H_2O} = \frac{2\pi c\sqrt{\epsilon_{m,\infty} + 2\epsilon_{H_2O}}}{\omega_P} \tag{S.3}$$

We can then rewrite $\lambda_{H_2O}$ in terms of $\lambda_{PVP}$:

$$\lambda_{H_2O} = \lambda_{PVP}\frac{\sqrt{\epsilon_{m,\infty} + 2\epsilon_{H_2O}}}{\sqrt{\epsilon_{m,\infty} + 2\epsilon_{PVP}}} \tag{S.4}$$

The plasmonic peak of nanostars coated in PVP is measured experimentally to be at 740nm. For gold $\epsilon_{m,\infty} = 7.9$,[6] for water $\epsilon_{H_2O} = 1.77$[7] and for PVP $\epsilon_{PVP} = 2.33$.[8]

Plugging these values in **Eq. (S.4)** yields $\lambda_{H_2O} = 706$ nm, which aligns almost perfectly with the position of the peak of Archon1[5] and could potentially correspond to the condition where the nanostars are immobilized on the glass substrates.

#### Estimation of the quantum yield enhancement of Cy5

To estimate quantitatively the fluorescence enhancement of Cy5 when bound to a nanostar via bBSA (**Fig. 3c** of the main manuscript), we calculated the average field enhancement in a ring at 13 nm from the surface to better represent a realistic effect (ring thickness 1 nm, 13 nm away from the surface).

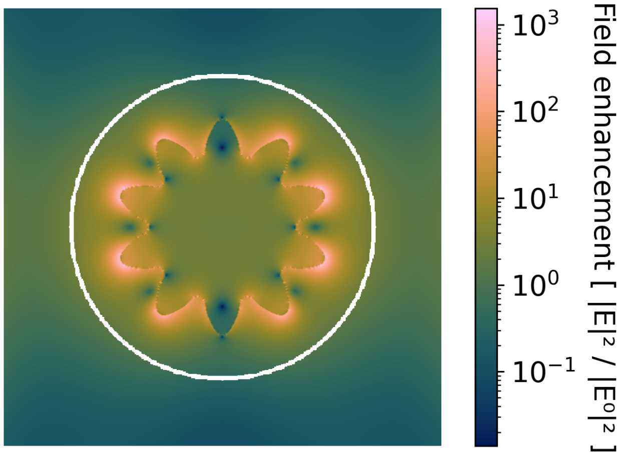


**Figure S10:** In white, the anular region around the nanostar where the field enhancement is averaged. Ring thickness 1 nm.

The average value in the ring is 1.78 (**Fig. S10**). Then, from a reference[9] we took $Y = 0.30$ and $\tau = 2.33$ ns and calculated:

$$\Gamma = \frac{Y}{\tau} = 0.129 \text{ ns}^{-1} \tag{S.5}$$

$$\Gamma_{nr} = \frac{1-Y}{\tau} = 0.300 \text{ ns}^{-1} \tag{S.6}$$

Given that the main effect is the enhancement of the radiative decay rate $\Gamma$, we can calculate the enhanced quantum yield using **Eq. (1)** (main manuscript):

$$Y_{enh} = \frac{\frac{|E|^2}{|E^0|^2} \cdot \Gamma}{\frac{|E|^2}{|E^0|^2} \cdot \Gamma + \Gamma_{nr}} = \frac{1.78 \cdot \Gamma}{1.78 \cdot \Gamma + \Gamma_{nr}} = 0.434 \tag{S.7}$$

the relative increase in the quantum yield is 45%, exactly what we measured.

#### Estimation of the glycocalyx spacing effect

Using the plasmonic ruler as calculated in the main manuscript, we can estimate the spacing due to the glycocalyx by comparing the field enhancement above and below the HEK cells. We use the quantum yield of Q6a $Y = 0.0367$.[10]

We know that:

$$Y = \frac{\Gamma}{\Gamma + \Gamma_{nr}} \Rightarrow \Gamma = \frac{Y}{1-Y}\Gamma_{nr} \tag{S.8}$$

and from our experiments:

$$Y_{below} = 1.69Y = 0.062 \tag{S.9}$$

$$Y_{above} = 1.28Y = 0.047 \tag{S.10}$$

The enhanced quantum yield due to field enhancement $\alpha$ is:

$$Y^* = \frac{\alpha\Gamma}{\alpha\Gamma + \Gamma_{nr}} = \frac{\alpha Y}{\alpha Y + (1-Y)} \tag{S.11}$$

Therefore we can express the field enhancement as:

$$\alpha = \frac{Y^*}{1-Y^*} \cdot \frac{1-Y}{Y} \tag{S.12}$$

For the case of nanostars embedded in the fibronectin coating, we get:

$$\alpha_{below} = 1.73 \tag{S.13}$$

which corresponds, according to the plasmonic ruler, to 13.5 nm. For the case of nanostars dropped above the cells, we get:

$$\alpha_{above} = 1.29 \tag{S.14}$$

which corresponds, according to the plasmonic ruler, to 16 nm. Knowing that the fibronectin is 6 nm thick (**Fig. 2**), the thickness of the glycocalyx can be estimated to be:

$$\Delta d = (16 - (13.5 - 6))nm = 8.5nm \tag{S.15}$$

### Photocycle fitting

The quantum yield of the fluorescent state is:

$$Y = \frac{\Gamma}{\Gamma + \Gamma_{nr}} \tag{S.16}$$

The enhanced quantum yield due to field enhancement $\alpha$ is:

$$Y^* = \frac{\alpha\Gamma}{\alpha\Gamma + \Gamma_{nr}} \approx \alpha Y \tag{S.17}$$

The approximation is valid for small $\Gamma/\Gamma_{nr}$. The measured fluorescence is the product of some efficiency factor (system dependent) $\eta$, the occupancy of state $Q$ and the quantum yield $Y$:

$$F = \eta Q Y \tag{S.18}$$

$$F^* = \eta Q^* Y^* = \eta Q^* \alpha Y \tag{S.19}$$

Brightness enhancement:

$$\frac{F_l^* - F_l}{F_l} = \frac{\eta Q_l^* \alpha Y - \eta Q_l Y}{\eta Q_l Y} = \alpha \frac{Q_l^*}{Q_l} - 1 \tag{S.20}$$

Voltage sensitivity (control):

$$\frac{F_h - F_l}{F_l} = \frac{\eta Q_h Y - \eta Q_l Y}{\eta Q_l Y} = \frac{Q_h}{Q_l} - 1 \tag{S.21}$$

Voltage sensitivity (enhanced):

$$\frac{F_h^* - F_l^*}{F_l^*} = \frac{\eta Q_h^* \alpha Y - \eta Q_l^* \alpha Y}{\eta Q_l^* \alpha Y} = \frac{Q_h^*}{Q_l^*} - 1 \tag{S.22}$$

These quantities so defined can be used to directly compare the features between measurements (where $F$ is known, but not the photocycle state population $Q$) and the simulations (where $Q$ is known, but not the quantum yield $Y$ or the efficiency factor $\eta$).

By implementing these criteria in a differential evolution algorithm (code available on GitHub[1]), we obtained a set of parameters and results for each case (**Table S1** and **S2**, simulated traces in **Fig. S11**). Note that the measured time constant of the enhanced response was likely limited by the camera response time, rather than the protein kinetics.

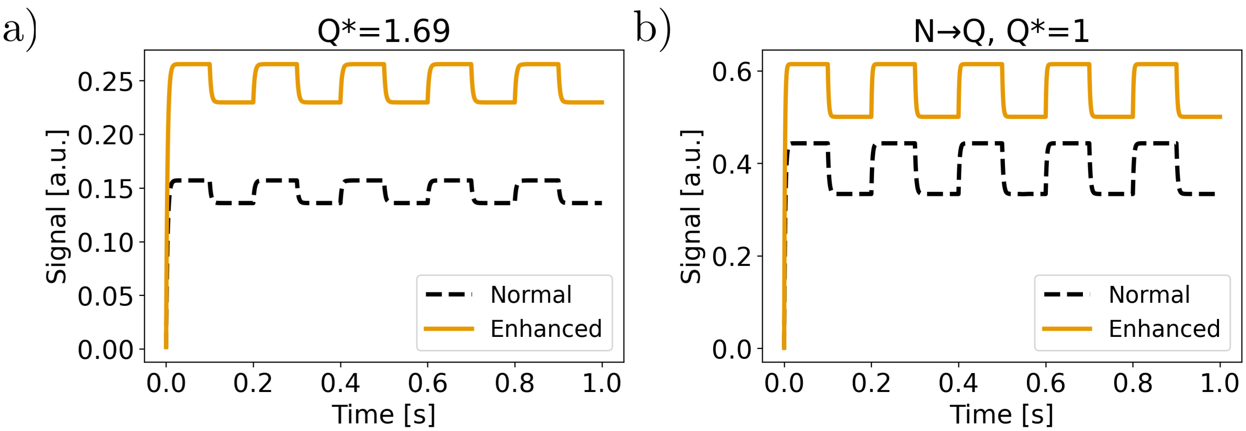


**Figure S11:** Numerically simulated traces for the apparent occupancy of the Q-state of QuasAr6a under the influence of a square wave in membrane potential (−70 mV to +30 mV). **a,** Plasmonic enhancement is assumed to enhance the Q-state only. **b,** Plasmonic enhancement is assumed to enhance $k_3$ only.

**Table S1:** Best parameters from genetic algorithm fitting of the experimental data.

| **Parameter** | | **Q*** | **N →Q** | **N →Q, Q*** |
|---|---|---|---|---|
| $k_1$ [s$^{-1}$] | | 12338 | 91991 | 99755 |
| $k_2$=$k_{2m}V + k_{2q}$ | $k_{2m}$ [V$^{-1}$s$^{-1}$] | 219915 | 284200 | 206712 |
| | $k_{2q}$ [s$^{-1}$] | 30695 | 29085 | 19088 |
| $k_3$ [s$^{-1}$] | Control | 100 | 252 | 443 |
| | Enhanced | 100 | 505 | 1184 |
| $k_4$ [s$^{-1}$] | | 261 | 232 | 119 |
| $k_5$ [s$^{-1}$] | | 9851 | 9800 | 9723 |
| $\left\|\vec{E}\right\|^2 / \left\|\vec{E}^0\right\|^2$ [a.u.] | | 1.69 | 1 | 1.34 |

**Table S2:** Comparison of the features and quantification of the Huber loss for experimental and simulated conditions.

| **Output** | | **Observed results** | **Q*** | **N →Q** | **N →Q, Q*** |
|---|---|---|---|---|---|
| Brightness Enhancement [a.u.] | | 0.69 | 0.69 | 0.50 | 0.88 |
| $\frac{\Delta F}{F}$ [a.u.] | Control | 0.37 | 0.16 | 0.33 | 0.33 |
| | Enhanced | 0.11 | 0.16 | 0.23 | 0.15 |
| $k_{up}$ [s$^{-1}$] | Control | 230 | 299 | 412 | 414 |
| | Enhanced | 1300 | 299 | 598 | 900 |
| $k_{down}$ [s$^{-1}$] | Control | 270 | 303 | 343 | 250 |
| | Enhanced | 930 | 303 | 455 | 459 |
| Huber loss [a.u.] | | | 14.8 | 11.6 | 8.8 |
